## Supporting Information for "Predictability and parallelism in the contemporary evolution of hybrid genomes"

**Supporting Information 1. Summary of past research on the ecological and mate choice environment in hybrid populations**

Hybrid populations formed between *X. birchmanni* and *X. malinche* have been better studied in an ecological context than hybrid populations formed between *X. birchmanni* and *X. cortezi*. *X. birchmanni* and *X. malinche* are adapted to distinct thermal environments, with *X. malinche* being limited to cool, high elevation sites [1,2]. Natural hybrids are found at a range of elevations, spanning *X. malinche* typical to *X. birchmanni* typical elevations, and tend to have intermediate thermal tolerance [2]. Importantly, genome-wide ancestry in *X. birchmanni* × *X. malinche* hybrid populations does not track expectations given the elevations at which populations are found. For example, the Tlatemaco population is majority *X. malinche* in its genome-wide ancestry (Fig. 1) but is found at a much lower elevation (~400 meters) than two populations with majority *X. birchmanni* ancestry, Totonicapa (upstream of Acuapa, ~700 meters) and Aguazarca (~900 meters; [1,3]). Much less is known about the ecological differentiation between *X. birchmanni* and *X. cortezi*. *X. cortezi* tends to be found at lower elevations than *X. birchmanni* and existing data indicates *X. cortezi* populations generally experience higher temperatures than *X. birchmanni* as well as greater seasonality [4]. Both *X. birchmanni* × *X. cortezi* hybrid populations are found at *X. cortezi*-typical elevations (~160-170 meters) and individuals in these populations derive the majority of their genomes from *X. cortezi*. Thus, connections between genome-wide ancestry and the ecological environments in which hybrids are found are not straightforward, and varying ecological conditions exist across distinct populations.

Similarly, the degree of assortative mating and differentiation in sexually selected traits varies dramatically across different replicate populations. Within *X. birchmanni* × *X. malinche* hybrid populations, strong ancestry assortative mating is observed at the Aguazarca population, with individuals of primarily *X. birchmanni* ancestry (Fig. 1) discriminating against individuals with high *X. malinche* ancestry [3], and vice versa. However, in two other hybrid populations, Totonicapa (upstream of Acuapa) and Tlatemaco, there is no evidence for deviations from random mating by ancestry [3]. Within both *X. birchmanni* × *X. cortezi* hybrid populations, we also see evidence for strong ancestry assortative mating [5]. Naively, this might lead to the prediction that similarities in local ancestry between Aguazarca and Huextetitla or Santa Cruz could be driven by assortative mating. However, initial data points to differences in the cues that are important for assortative mating in the two different hybrid population types. For example, in Aguazarca, loci underlying the sexually selected sword ornament have resisted introgression, whereas the opposite is true in in Huextetitla and Santa Cruz [5].

Together, the large variation observed in the ecological and mate choice environments across the three *X. birchmanni* × *X. malinche* and two *X. birchmanni* × *X. cortezi* hybrid populations indicate that these factors are unlikely to drive correlations in local ancestry across populations that are consistently observed (i.e. minor parent deserts that are shared across most populations). However, these factors may play an important role in explaining shared local ancestry between pairs of populations.

**Supporting Information 2. Principal Component Analysis of within populations variation in low coverage data from Santa Cruz and Huextetitla**

For the purposes of our analyses, we wanted to know whether the Santa Cruz and Huextetitla populations formed independently and whether they have been independent in their recent demographic history. We took advantage of the fact that our local ancestry data allows us to identify the locations of recombination events between haplotypes of different ancestry. These ancestry transitions reflect not only recombination events that occurred in the previous generation but ancestral recombination events that have been inherited in sampled individuals. Thus, the set of ancestry transitions found in an individual is one sample of the recombination history of a population. Distinct recombination histories in Huextetitla and Santa Cruz would support the conclusion that these populations have largely independent recent histories. Using the locations of ancestry transitions in both populations (see Methods), we performed a PCA on recombination event presence or absence across windows in R. Intriguingly, Huextetitla and Santa Cruz are well separated along PC2, which explains 1.5% of the variance in the distribution of recombination events (Fig. S3). PC1, which correlates with genome-wide ancestry ($R=0.93$, $p<10^{-42}$), explains 5% of the variance in the data. However, we noticed that there was some overlap between the two populations along PC2 (Fig. S3), potentially indicative of recent migration.

To further explore whether there is differentiation in the genetic composition of these populations we again used a principal component analysis approach. Since our population level data is low-coverage, we started with raw count data extracted from individual gvcf files generated with samtools and bcftools [6,7] for 50 randomly sampled individuals from Santa Cruz and all available individuals from Huextetitla. At sites with reads that supported more than one allele, we used binomial sampling to randomly select one read to represent the variant site for that individual and converted the output into plink format using a custom script (https://github.com/Schumerlab/Lab_shared_scripts/random_flip_samtools_low_coverage_vcf_for_PCA.pl). We removed non-biallelic sites and used the plink merge function to merge all individuals [8]. We used plink to remove sites with >50% missing data in our sample and with minor allele frequencies of less than 2%. After performing this filtering, we used the pca function in plink to perform principal component analysis for the first 20 principal components. Visualizing this analysis indicated that Santa Cruz and Huextetitla were clearly separated along PC2 (Fig. S3). PC1 and PC2 explain 7.3 and 3% of the variance in the data respectively. In addition, we previously reported substantial differences in the genetic diversity of *X. cortezi* ancestry tracts in the Huextetitla and Santa Cruz populations ([5]; Fig. S2). This suggests that SNP variation within ancestry tracts varies between Santa Cruz and Huextetitla, supporting some differences in contribution or retention of parental haplotypes from the source *X. cortezi* or *X. birchmanni* populations.

Together this data suggests that the Huextetitla and Santa Cruz hybrid populations have been independent in their recent evolutionary history, with some evidence for migration between them (Fig. S2). We also analyzed correlations in local ancestry across the Huextetitla and Santa Cruz hybrid populations (see Methods), reasoning that very high correlations in local ancestry could indicate that these populations have been connected historically. We found extremely strong correlations in local ancestry across the two populations (Fig. S3; Table S10). Partial correlation analyses indicated that some of this signal could be explained by shared recombination maps and shared locations of coding and conserved basepairs. However, substantial signal remains even after accounting for these features (100 kb windows: $\rho = 0.72$, $p<10^{-300}$; see Table S12, S13 for other window sizes).

Cross-population correlations in ancestry are unexpected in the absence of shared sources of selection on hybrids (see Supporting Information 3,6) and/or shared population history. We used simulations to investigate whether shared sources of selection could plausibly generate the magnitude to cross-population correlations in ancestry that we observed. To do so, we simulated a hybrid population formed between two species, each with 24 chromosomes that were 25 Mb in length. We performed simulations using the forward-time simulation program admix'em [9] which is designed to simulate admixture. We used the local recombination map for the 24 chromosomes from *X. birchmanni* to determine recombination probabilities and tracked ancestry at 24,000 markers. Based on estimates from previous work on the Santa Cruz and Huextetitla populations [5], we implemented a pulse of admixture 150 generations ago, a hybrid population size of 3,000 diploid individuals, and admixture proportions of 85% parent 1 and 15% parent 2. We simulated two replicate hybrid populations that formed independently and sampled 242 individuals from one population and 12 from the other. To implement selection on hybrids in simulations, we modeled selection against Dobzhansky-Muller hybrid incompatibilities, as these types of interactions appear to be common in *Xiphophorus* species [10–12]. For each selection scenario we performed 25 replicate simulations. We simulated the following scenarios:

1. Selection on five randomly placed pairs of Dobzhansky-Muller hybrid incompatibilities, each with a selection coefficient of 0.2 and dominance coefficient of 0.5
2. Selection on thirty randomly placed pairs of Dobzhansky-Muller hybrid incompatibilities, each with a selection coefficient of 0.1 and dominance coefficient of 0.5

Our simulations indicated that shared sites under selection can drive substantial correlations in local ancestry across independently formed hybrid populations. For scenario 1 we observed cross population correlations in local ancestry up to $\rho = 0.41$ (median $\rho$ across simulations = 0.19; summarized in 250 kb windows). For scenario 2 we observed cross population correlations in local ancestry up to $\rho = 0.6$ (median $\rho = 0.43$).

Although both sets of simulations induced substantial correlations in local ancestry across populations, these correlations were much weaker than those observed in the real data. Even in scenario 2 where we implemented strong selection on many loci, cross-population correlations did not approach the correlation coefficients we observe in the real data for Huextetitla and Santa Cruz. Since even models of extremely strong selection are not sufficient to generate the cross-population ancestry correlations we observe in Huextetitla and Santa Cruz (Table S10), we conclude that Huextetitla and Santa Cruz are unlikely to be truly independent hybridization events between *X. birchmanni* × *X. cortezi*. We thus focus our analyses in the main text on the Santa Cruz hybrid population, where we have a much larger sample of hybrids (N=242).

**Supporting Information 3. Simulations fitting the demographic history of Santa Cruz and Acuapa**

In addition to selection on hybrids, the demographic history of hybrid populations can drive variation in ancestry across the genome. For example, ancestry tracts can be fixed by genetic drift even in young hybrid populations if population sizes are sufficiently small.

*Inference of demographic history using an ABC approach*

We wanted to evaluate whether patterns of fixation or loss of minor parent ancestry could be explained by the demographic history of the focal hybrid populations. To do so, we used a simulation-based approach. For the purposes of simulations, we focus on modeling one *X. birchmanni* × *X. cortezi* and one *X. birchmanni* × *X. malinche* hybrid population. We previously used an approximate Bayesian computation approach in SLiM [13] to explore the demographic history of the Santa Cruz hybrid population [5]. We build upon this approach here and also apply it to infer the demographic history of the Acuapa hybrid population.

The overall structure of the simulations implemented here were as follows: we drew population demographic parameters from uniform prior distributions, performed simulations in SLiM [14], generated summary statistics from these simulations, and compared them to the real data to accept or reject each simulation. Because we focus on ancestry in hybrids rather than population genetic summaries, we initialized two parental populations for the purposes of each simulation and immediately formed a hybrid population between them. We simulated a 25 Mb chromosome with recombination rates matching those observed on chromosome 2 of *X. birchmanni*. We used tree sequence recording in SLiM to track ancestry tracts in each individual in the simulated population [14]. Parameters for the time since initial admixture (0-300 generations), admixture proportion (0.6-1 of the genome derived from the major parent species), and hybrid population size (100-8,000 diploid individuals), were drawn from uniform prior distributions. For migration rate from each parental species, we drew from a log uniform prior distribution of 0-5%. We chose to use a log uniform prior distribution because the absence of parental individuals and early generation hybrid individuals in population samples suggested that migration rates are low.

For each simulation, we sampled the number of individuals that matched the sampling effort in our empirical datasets (N=97 for simulations of Acuapa and N= 242 for Santa Cruz) and summarized average ancestry, the coefficient of variation in chromosome-wide ancestry, and the median length of ancestry tracts derived from the minor parent species. We accepted simulations where the simulated values for all summary statistic fell within ±5% of the focal values. We also repeated this procedure with local variation in ancestry as a summary statistic, using the coefficient of variation of ancestry in 250 kb windows (see below).

As a proof of principle, we first evaluated how well this procedure performed with known parameters. Specifically, we randomly sampled a simulation with known input parameters and treated the summary statistics from that simulation as if they were the real data. Using these summary statistics, we collected all other simulations that passed our criteria for acceptance, requiring that each randomly sampled parameter set yielded at least 500 accepted simulations. For parameter sets with more than 500 accepted simulations, we randomly sampled 500 of them. We then calculated the maximum a posteriori or MAP estimate and 95% quantile range of the posterior distribution. Next, we asked how well the posterior distributions from this step captured the known input parameters for the focal simulation. To do so, for each parameter, we recorded the difference between the MAP estimate and the true parameter value, as well as whether the true value of the parameter fell in the 95% quantile of the posterior distribution. We repeated this procedure for 5,000 randomly sampled pairs of parameter sets and simulation summary statistics.

Based on this analysis, we found that MAP estimates from our ABC procedure frequently fell close to the true simulated parameter values (Fig. S28) and that >90% of the time, the true parameter value fell within the 95% quantile of the posterior distribution.

For the real data, we followed the same procedure, accepting simulations where the simulated values for all summary statistic fell within ±5% of the observed values for the real data. We performed simulations until 500 parameter sets had been accepted. We recovered well-resolved posterior distributions for all parameters for both populations (Fig. S4). We recorded each set of accepted parameters for each population for use in subsequent simulations.

*Results using a summary statistic of local ancestry*

We used two sets of summary statistics to infer demographic parameters from simulated data. In the first approach, we use summaries of average ancestry, and the coefficient of variation in chromosome-wide ancestry, and the median length of ancestry tracts derived from the minor parent species. In the second approach, we use a summary of variation in local ancestry rather than chromosome-wide ancestry (the coefficient of variation in ancestry summarized in 250 kb windows). With this second approach, we observed a marked reduction in the probability of a simulation being accepted (~10X) and were not able to accept a sufficient number of simulations for either population to proceed with the approach (i.e. fewer than 100 accepted simulations for either population out of ~1 million simulated parameter sets).

Visualizing the accepted simulations, this approach yielded similar estimates of most demographic parameters (Fig. S4), but resulted in a narrower posterior distribution for hybrid population size. Interestingly, this approach yielded estimates of hybrid population sizes that were very low, and in the case of Santa Cruz, was close to the number of individuals sampled for this project. Because we have never recaptured a previously fin-clipped individual in either hybrid population, we believe that these population size estimates are unrealistically low, and likely reflect the fact that increased genetic drift is required to describe local ancestry variation in the absence of selection. However, we view these neutral simulations as an important null hypothesis in investigating local ancestry variation, as described below.

*Simulation of variation in local ancestry given estimated demographic parameters*

Using inferred parameters for the demographic history of the Acuapa (*X. birchmanni* × *X. malinche* hybrid population) and Santa Cruz population (*X. birchmanni* × *X. cortezi* hybrid population), we proceeded to simulate local ancestry in hybrid populations under these parameters. We performed 100 replicate simulations for each hybrid population. We used the admix'em simulation framework described in Supporting Information 2. For each of these simulations we randomly drew parameter sets from the posterior distribution of ABC simulations described above (Fig. S4). When simulations finished we summarized average ancestry from a random sample of 242 and 100 individuals for the Santa Cruz and Acuapa simulations respectively. We performed analyses in 10, 100, and 250 kb windows as we had done for the real data. We repeated this procedure with accepted ABC simulations that incorporated local ancestry variation (see previous section and Fig. S4), except that we only performed 50 replicate simulations.

While average minor parent ancestry in simulations matches that observed in our real data for both Acuapa and Santa Cruz, variation in local ancestry along the genome greatly exceeds expectations from simulations in both populations and both simulation scenarios (Fig. S5). This suggests that other drivers of local ancestry variation besides demographic history, such as selection, may be required to explain the observed data. Indeed, past work has suggested a substantial role of selection against hybrid incompatibilities in driving local ancestry variation in *X. birchmanni* × *X. malinche* hybrid populations [15].

We also asked about cross-correlations in ancestry in the two types of simulated hybrid populations under these neutral demographic scenarios. As expected, we did not observe correlations in minor parent ancestry across simulations of the two types of hybrid populations neutral simulations (Fig. S10).

**Supporting Information 4. Expectation that local recombination rates and locations of functional basepairs are conserved between *X. birchmanni* and *X. cortezi***

In the main text, we use a linkage disequilibrium (LD) based recombination map that we previously generated for *X. birchmanni* to analyze correlations between minor parent ancestry, recombination rate, and other genomic features of interest [15]. Our approach thus assumes that local recombination rates are conserved between *X. birchmanni* and *X. cortezi*. Past work has shown that swordtails and other percomorph fish carry a truncated version of the PRDM9 gene [16], the gene responsible for regulating the locations of recombination hotspots in mammals. This truncated gene is not active in specifying the locations of recombination hotspots [15,16]. Importantly, species groups that lack an active PRDM9 exhibit strong conservation of the locations of recombination hotspots, even over deep evolutionary timescales [17,18]. Given that *X. birchmanni* and *X. cortezi* are close relatives, and we previously found no evidence for divergence in recombination maps between *X. birchmanni* and *X. malinche* [15], we are confident in using the *X. birchmanni* recombination map to analyze covariates with ancestry in *X. birchmanni* × *X. cortezi* hybrids. One important exception is at large inversions that distinguish the two species on chromosome 21 and 24 (Fig. S18) and smaller inversions elsewhere in the genome (see Methods). Excluding chromosomes 21 and 24 from our analysis had no impact on our results (Table S2).

To further confirm our expectation that *X. birchmanni* and *X. cortezi* share fine-scale recombination maps, we evaluated evidence for biased gene conversion in *X. cortezi* at hotspots and matched coldspots that were first identified in *X. birchmanni*. Specifically, we identified 5 kb regions from the *X. birchmanni* LD map that had an estimated heat greater than 10X the background rate (determined from the flanking 20 kb windows with a 5 kb spacer). We identified matched coldspots by searching for 5 kb regions that were between 0.9 to 1.1X the background rate but had GC content within 10% of that observed in the focal window. We repeated this procedure until each hotspot had a matched coldspot. We then calculated GC* in each hotspot and matched coldspot using the relationship:

$$GC^* = \frac{\frac{AT \rightarrow GC}{ancAT}}{\frac{AT \rightarrow GC}{ancAT} + \frac{GC \rightarrow AT}{ancGC}}$$

Where *ancAT* and *ancGC* are the counts of A/T or G/C in the inferred ancestral sequence and AT → GC and GC → AT represent the number of substitutions of each type in the focal lineage relative to the inferred ancestry sequence. To determine *ancAT* and *ancGC*, we used a previously inferred ancestral sequence for northern swordtails [15], which was generated using the prequel option in phastcons [19]. We restricted analysis to sites that were assigned a greater than 0.9 probability of a given basepair representing the ancestral state. We compared GC* in each hotspot and matched coldspot. The results of this analysis are shown in Fig. S6.

Since we similarly use the coordinates of functional elements based on the *X. birchmanni* reference genome for our analyses of covariates with minor parent ancestry in the main text, we also evaluated the degree of synteny across *X. birchmanni*, *X. malinche*, and *X. cortezi*. Across the genome, the three species differ by three large inversions on chromosomes 17, 21, and 24 as well as 23 smaller inversions with an average length of 146 kb (7 derived in *X. cortezi*, 8 derived in *X. birchmanni* and 8 derived in *X. malinche*). To ensure that these larger and smaller scale breakdowns in synteny were not impacting our results, we repeated our analyses of the correlations between coding and conserved basepairs and minor parent ancestry, excluding windows that overlapped with inversions. Our results were qualitatively unchanged (Table S5).

**Supporting Information 5. Evaluating the signal of lower minor parent ancestry in regions of the genome with more coding substitutions between species**

We detected a modest correlation between the frequency of minor parent ancestry and the density of coding substitutions. Although one might conclude from this data that more differentiated regions of the genome resist introgression between species, an equally likely explanation is that improved power to infer ancestry in differentiated regions drives this signal.

To evaluate this possibility, we thinned ancestry informative markers in regions where the density of ancestry informative markers exceeded the median density genome wide (as described in *Local ancestry inference in X. birchmanni* × *X. cortezi hybrids*) and re-inferred ancestry. We found that the relationship between minor parent ancestry and coding substitutions remained (Table S6), indicating that it is not an artefact of power to infer ancestry.

Somewhat surprisingly, we found that the effect size is not different between synonymous or nonsynonymous changes in the cDNA sequence (Table S7). We evaluated this in more detail by matching 0.1 cM windows for the total number of total coding substitutions and comparing ancestry between them. Specifically, we identified 0.1 cM windows in the upper 25% quantile of the number of nonsynonymous substitutions between *X. cortezi* and *X. birchmanni*. We then identified a matched window that contained no nonsynonymous substitutions but a similar number of synonymous substitutions (within 20%). Comparing the ancestry distributions between these sets of matched windows indicated that they did not differ in minor parent ancestry (Fig. S8). These results support the idea that the minor parent ancestry patterns around coding substitutions are driven more by the locations of coding basepairs rather than the identity of nearby substitutions. Indeed, given that ancestry tracts are so long in early generation hybrids we may not expect to see differences in ancestry around individual functionally important sites, unlike what has been reported in more ancient hybridization events [20].

**Supporting Information 6. Simulations to explore mechanisms of selection on hybrids**

*Observed correlations with minor parent ancestry within populations*

We observe positive correlations between minor parent ancestry and the local recombination rate in both Huextetitla and Santa Cruz hybrid populations, and we previously reported similar correlations in all three *X. birchmanni* × *X. malinche* hybrid populations ([15]; Acuapa/Totonicapa, Aguazarca, and Tlatemaco). As we demonstrated in simulations described in Supporting Information 3, these correlations between local ancestry and the recombination rate are unexpected under scenarios of neutral admixture (Fig. S10; Fig. S29), even when matching the demographic history of the Santa Cruz and Acuapa hybrid populations (Supporting Information 3).

Given that neutral admixture does not typically generate the correlations between ancestry and recombination rate that we observe, we next asked whether selection could drive the correlations we observe. While we cannot explore all possible modes of selection on hybrids for practical reasons, we perform simulations of two general cases that appear to commonly impact hybrid fitness [21]. In the first set of simulations, we implement incompatibility selection on hybrids. In the second set of simulations, we treat variants from one of the parental species as globally deleterious, but weakly so.

We again drew from the posterior distributions of accepted ABC simulations to select demographic parameters for each simulation. First, we performed simulations where we randomly placed 20 pairs of hybrid incompatibilities along the genome, each with $s$=0.1 and $h$=0.5. Next, we performed simulations where there was global selection against ancestry derived from one of the parent species (arbitrarily *X. birchmanni* in these simulations). This was implemented by drawing 100 random sites where *X. birchmanni* ancestry was disfavored with $s$=0.01 and $h$=0.5, such that the total strength of selection on $F_1$ hybrids was the same across simulations. We performed 100 replicate simulations for each scenario.

Both sets of simulations resulted in significant correlations between local ancestry and the local recombination rate (Fig. S10; Fig. S29). The directions of these correlations further allowed us to evaluate likely sources of selection driving the patterns that we observe in the real data. Simulations of selection against hybrid incompatibilities consistently resulted in positive correlations between minor parent ancestry and the local recombination rate (summarized in 250 kb windows; Fig. S10, Fig. S29). In contrast, simulations of global selection against one parent species resulted in a positive correlation between recombination rate and minor parent ancestry in one set of simulations and a negative correlation between recombination rate and minor parent ancestry in the other set of simulations (Fig. S10; Fig. S29).

In the real data, the observed patterns are consistent with selection against minor parent ancestry in all hybrid populations (Fig. 2, Table S1; [15]). Moreover, we observe a positive correlation between minor parent ancestry and recombination rate regardless of the identity of the minor parent species (i.e. *X. birchmanni*, *X. cortezi*, or *X. malinche*). These results indicate that selection against a particular parental ancestry type (e.g. as under a hybridization load model), cannot explain the patterns in our data. Similarly, we suspect that ecological selection or sexual selection on hybrids could not drive these global patterns, given what is know about these factors across population (see Supporting Information 1 for more details).

What mechanisms of selection could drive the patterns of minor parent ancestry observed in the real data? Hybrid incompatibility selection, as we simulate above, drives purging of minor parent ancestry at loci involved in incompatibilities. Since this mechanism can generate the patterns observed in our data and there are many documented cases of hybrid incompatibilities in *Xiphophorus* [10–12], this may be the most likely mechanism of selection generating the patterns that we observe. However, any mechanism of selection that drives purging of minor parent ancestry across the genome could generate the patterns we observe. Two other possible mechanisms that have been understudied in empirical literature include stabilizing selection on quantitative traits in hybrids or selection under Fisher's geometric model, where removal of minor parent ancestry shifts hybrids closer to the major parent's phenotypic optimum [21].

*Observed correlations in minor parent ancestry between populations*

In addition to correlations between minor parent ancestry and local recombination rate within hybrid populations, we also observe correlations in local ancestry *between* the two types of hybrid populations (Table S10, S12). Even after accounting for local recombination rate and the number of linked coding and conserved basepairs using a partial correlation approach, we still observe substantial correlations in minor parent ancestry across some pairs of populations (e.g. in 250 kb windows for Santa Cruz and Acuapa: $\rho = 0.16$, $p=10^{-16}$; Table S12). This suggests that genetic architecture alone cannot explain the correlations in local ancestry between the two types of hybrid populations.

We used simulations to explore how shared selection could generate the correlations in minor parent ancestry we observe across different hybrid population types. These simulations are similar to those described in Supporting Information 2 but incorporate demographic history of the Santa Cruz and Acuapa hybrid populations as described in Supporting Information 3. For simulations of selection on hybrids there are many unknown parameters. Because of this, we simply asked whether randomly placing selected sites in the same locations in the two simulated hybrid populations could generate patterns qualitatively similar to the cross-population correlations in local ancestry that we observe. We performed 100 replicate simulations under two scenarios of selection:

1. Selection against minor parent ancestry at 5 randomly placed pairs of sites along the genome ($s$=0.1, $h$=0.5).
2. Selection against minor parent ancestry at 20 randomly placed pairs of sites along the genome ($s$=0.1, $h$=0.5).

As with the real data, we performed partial correlation analysis, asking if there were correlations in minor parent ancestry between simulated Santa Cruz and Acuapa populations after accounting for local variation in recombination rate. We did not account for the locations of coding (or conserved) basepairs in this analysis since we did not use an exon map to place selected sites in simulations.

We found that both selection scenarios resulted in significant positive correlations in ancestry across the two hybrid population types (Fig. S10; at $p < 0.05$ for 91% of simulations for scenario 1, 97% for scenario 2). Interestingly, the magnitude of cross-population correlations produced in the second simulation scenario is more consistent with that observed in the real data between the Santa Cruz and Acuapa hybrid populations (mean $\rho$ scenario 1: 0.11, mean $\rho$ scenario 2: 0.24). This could indicate that a large number of shared sites under selection are required to generate the cross-population correlations that we observe. However, we caution that there are many variables we are not modeling in these simple simulations, such as variation in the strength of selection on individual loci and loci that are under selection in one hybrid population type and not the other. While modeling this complex parameter space is beyond the scope of our analysis here, it will be an exciting direction for future work.

*Reduced correlation in minor parent ancestry in comparisons involving the Tlatemaco population*

In our analysis of the empirical data, one replicate *X. birchmanni* × *X. malinche* hybrid population, the Tlatemaco population, is an exception to the global pattern of cross population correlations in local ancestry. Genome-wide, minor parent ancestry in Tlatemaco is poorly predictive of minor parent ancestry in both other *X. birchmanni* × *X. malinche* populations and in *X. birchmanni* × *X. cortezi* hybrid populations (Fig. 3; Table S10, S12). Naively, since *X. birchmanni* is the minor parent species in both Tlatemaco and in *X. birchmanni* × *X. cortezi* hybrid populations, we might expect the opposite pattern (stronger cross-correlation between Tlatemaco, Santa Cruz, and Huextetitla). We speculate that some of this break down in repeatability could be attributable to the fact that hybrids in Tlatemaco derive the majority of their genome from *X. malinche* (Fig. 1), which experienced a strong and persistent bottleneck relative to *X. birchmanni* and *X. cortezi* (Fig. S1, [5,15]). As a result, some *X. malinche*-derived haplotypes may have higher genetic load and be globally deleterious [22].

Even if increased genetic load in *X. malinche* haplotypes were driving lower cross-population correlations with other hybrid populations genome-wide, we would still expect to see reduced minor parent ancestry in Tlatemaco in regions of the genome where minor parent ancestry is strongly deleterious (i.e. near hybrid incompatibilities). Indeed, when we evaluate ancestry near segregation distorters in Tlatemaco, we see a reduction in minor parent ancestry closer to these strongly selected sites (Fig. S30, $\rho$= -0.12, $p<0.0002$; analysis conducted in 10 kb windows that fall within 500 kb of a segregation distorter). We also still observed enrichment in shared minor parent deserts between Tlatemaco and *X. birchmanni* × *X. cortezi* hybrid populations (Fig. 3), consistent with the idea that there are several regions genome-wide that are under shared selection between Tlatemaco and the other hybrid populations studied here.

### Supporting Information 7. Power to detect shared minor parent deserts

In the main text, we perform a scan for shared minor parent deserts as part of our analysis of patterns of ancestry in *X. birchmanni* × *X. cortezi* and *X. birchmanni* × *X. malinche* hybrid populations. To explore what selection coefficients we expect to have power to detect in this analysis, we used a simulation-based approach similar to those described above.

For each simulation, we again drew demographic parameters randomly from the accepted posterior distributions for the Santa Cruz and Acuapa populations separately (as described in Supporting Information 3,6) and performed simulations using admix'em [9]. For each pair of simulations, we randomly selected one site across the 24 chromosomes to be under selection in hybrid populations. Since we do not know the mechanisms of selection driving the emergence of shared deserts, we simply implemented negative selection against minor parent ancestry at these randomly selected sites.

Next, we allowed selection to occur at the same sites independently in the two hybrid populations and then converted the ancestry output from 242 individuals in the simulated Santa Cruz population and 97 individuals in the simulated Acuapa population to the same format as the hard-calls in our real data. We averaged ancestry by site in the simulated Santa Cruz population data and used this simulated data to identify minor parent ancestry deserts as we had in the real data. We used the same criteria as described in the Methods to determine if the ancestry desert was shared across the simulated Santa Cruz and Acuapa populations. We performed simulations using a range of selection coefficients ($s$=0.025, 0.05, 0.1). For each selection coefficient, we performed 100 replicate simulations.

We summarized the proportion of shared ancestry deserts that were identified at sites that were truly under selection in our simulations. Our results indicate that we have good power to detect shared minor parent ancestry deserts across a range of selection coefficients, and that power generally increased as the strength of selection increased (*s*=0.025 – 41% detected, *s*=0.05 – 63% detected, *s*=0.1 – 74% detected).

**Supporting Information 8. Evaluating possible technical artifacts and other drivers of minor parent ancestry deserts and islands**

*Evaluating the impact of spatial autocorrelation in ancestry on the number of minor parent deserts and islands*

In the main text, we evaluate the probability that minor parent deserts and islands are shared across populations by chance by randomly sampling windows of a given genetic length in each population (see Methods), and asking how frequently ancestry outliers co-occur across populations by chance. While this approach has the advantage of being simple and thus easily interpretable, it ignores an important problem generated by autocorrelation in ancestry along the genome [23]. For example, one large ancestry desert that was shared across populations would be overcounted using window-based approaches, and the method we use to generate null expectations for ancestry sharing would not correct for this.

To evaluate the impact of autocorrelation in ancestry between adjacent windows in our estimation of the number of minor parent ancestry deserts and islands, we performed additional analyses. First, we used a correction described in Hahn 2006 [23] to account for autocorrelation in our estimates of the number of ancestry deserts and islands. This approach models the chromosome as a one dimensional space, regresses spatially ordered subsequent measurements on eachother, and uses the strength of autocorrelation to estimate the number of effective samples (i.e. in our case, deserts or islands). We applied this correction [23] to minor parent islands and deserts found on the same chromosome in the Santa Cruz hybrid population (where islands and deserts were initially identified, see Methods). We found that the estimated effective number of deserts and islands was similar after this correction (81 versus 75 and 76 versus 63, respectively), suggesting that with the approach we used to delineate these regions our estimates are not strongly inflated by spatial autocorrelation in ancestry.

While the above analysis reassuringly suggests that we are not overcounting minor parent islands and deserts in our focal dataset, it does not account for issues related to autocorrelation in our null datasets. Specifically, by bootstrapping windows to estimate the probability that ancestry outliers are shared across populations by chance, we disrupt spatial autocorrelation in ancestry, and this could lead to underestimation of the false positive rate. As a second approach to estimating the expected overlap in ancestry outliers between populations by chance that preserves the ancestry correlation structure of the real data, we used a label shifting approach to generate our null expectations. We shifted the observed minor parent ancestry values (summarized in 0.05 cM windows) in *X. birchmanni* × *X. malinche* populations by sliding windows by 12.5 cM and then comparing these shifted values to the observed values in Santa Cruz. This allowed us to compare overlap in minor parent ancestry outliers between Santa Cruz and permuted *X. birchmanni* × *X. malinche* datasets, while preserving local ancestry structure in the permuted datasets. For each shifted dataset, we asked whether any windows that were major or minor parent ancestry outliers in the shifted data overlapped with the ancestry deserts identified in Santa Cruz (using the same criteria as in the real data). We repeated this procedure 130 times to fully tile the whole genome. Consistent with our initial approach, we found that few minor parent ancestry outliers in *X. birchmanni* × *X. malinche* hybrid populations overlapped minor parent deserts identified in the Santa Cruz hybrid population by chance. For minor parent islands the approach used mattered more, but we still observed an enrichment in most populations. The results of the shifted dataset approach are summarized in Fig. S12.

*Examining covariates with minor parent deserts and islands*

One common observation in other systems is that there is enrichment of minor parent deserts on certain chromosomes, particularly the sex chromosome. We evaluated whether there was an enrichment of deserts or islands on any one chromosome, relative to expectations based on their size. To do this we took the number of observed islands and deserts and randomly permuted their positions across the genome and counted how many randomly landed on each chromosome. We ran 1,000 permutations for the deserts and islands independently and compared the permuted distribution to the observed count on each chromosome. While we observe enrichment on some chromosomes, we do not identify an excess of minor parent deserts or islands on chromosome 21, the putative sex chromosome (see Results).

Genome-wide, we identified a much larger number of shared minor parent ancestry deserts and minor parent ancestry islands across natural hybrid populations than expected by chance (see Fig. S31 for a plot of all shared deserts and islands). One possibility is that these shared regions are simply correlated with other features of the genome that tend to be under selection on hybrids. While we account for recombination rate and the spatial structure of ancestry correlation in our identification of shared minor parent ancestry deserts and islands, we do not account for other features that may impact ancestry. For example, introgression of particularly gene dense regions may be selected against not because the same variants are deleterious in hybrids but because there is a larger target size in these regions.

To evaluate this, we compared the number of linked coding and conserved basepairs in the 0.05 cM window that contains the midpoint of the shared minor parent ancestry deserts and islands to a background set of all 0.05 cM windows in the genome. We found that these regions did not contain an excess of linked coding or conserved basepairs compared to other regions, suggesting that differences in ancestry are not driven by differences in target size (Fig. S14). We also separated shared minor parent deserts and islands into quantiles based on the number of linked coding and conserved basepairs, and plotted ancestry in these regions relative to 0.05 cM windows within the same coding and conserved basepair quantile (Fig. S32). Shared minor parent ancestry deserts and islands were still ancestry outliers when compared only to windows with a high density of coding and conserved basepairs.

In addition, we asked whether minor parent islands were still ancestry outliers when compared to regions of the genome that are likely to have unusually low constraint. We used bedtools [24] to determine the distance of each 10 kb window to the nearest coding basepair in the *X. birchmanni* genome. We selected windows that were >100 kb away from the nearest coding basepair and plotted ancestry in these windows versus ancestry at minor parent islands. Results of this analysis are shown in Fig. S16.

Because the analyses described in the previous paragraphs make comparisons only to coding and conserved basepair density in 0.05 cM and 10 kb windows, we also wanted to perform comparisons that specifically captured the spatial structure of the minor parent deserts and islands. As such, we generated null datasets based on the footprint of minor parent deserts and islands in our empirical data. Specifically, for each minor parent desert or island, we randomly drew a chromosome and an interval equal to the observed length of that region. We repeated this procedure for each desert or island to generate a null dataset. We then performed this null dataset generation procedure 1,000 times and compared summary statistics from nulls to the real data. The results of this analysis are shown in Fig. S33. Based on this approach, we found that minor parent deserts showed a trend towards having fewer coding basepairs than expected by chance but did not contain an unusual number of conserved basepairs compared to matched null datasets. Minor parent islands showed a trend towards being enriched in coding basepairs based on this analysis approach and but harbored an average number of conserved basepairs. While not significant, the trends for minor parent deserts and islands for the number of coding basepairs actually ran counter to our naïve expectations. We initially predicted that if anything, minor parent deserts would harbor more coding basepairs (reflecting more constraint) and minor parent islands would harbor fewer coding basepairs (reflecting less constrain). That the trends we observe in this analysis run counter to those expectations again suggests that the architecture of the genome is unlikely to explain the existence of shared minor parent islands and deserts.

Another possible cause of shared minor parent ancestry deserts and islands is differences driven by technical artifacts. In order to explore this possibility, we evaluated a number of possible technical artifacts that, if systematically different, could impact the accuracy of ancestry calls and potentially cause spurious signals. We first evaluated the density of repetitive elements in minor parent ancestry deserts and islands, based on the locations of repetitive elements in the *X. birchmanni* genome [11]. High repetitive element content could cause difficulties in accurately inferring ancestry. However, we find no evidence of differences in the density of repetitive basepairs between minor parent ancestry deserts and islands and other regions of the genome (Fig. S15).

Another consideration is our power to accurately infer local ancestry. While simulations indicate that we have excellent power to infer local ancestry in *X. birchmanni* × *X. cortezi* hybrids given the density of ancestry informative sites [5], there may be subtle variations in power along the genome. We evaluated whether minor parent deserts or islands had an unusually high or low density of ancestry informative sites. We find no evidence for differences in power to infer ancestry in focal windows, compared to the genomic background (Fig. S15). We also confirmed that our conclusions were similar when evaluating the data based on ancestry probabilities inferred using a thinned dataset that was designed to reduce power differences in ancestry inference between different regions of the genome (see *Local ancestry inference in X. birchmanni* × *X. cortezi hybrids*). Based on this analysis, we found that some minor parent islands were inferred to have substantially different ancestry using the thinned data (greater than 10% difference in minor parent ancestry between the unthinned and thinned data; Fig. S13). We thus excluded these regions from further analysis.

*Alternative approaches to evaluate the likelihood of shared minor parent ancestry deserts and islands*

In the main text, we describe an approach for identifying shared minor parent deserts and islands. We then use two types of permutations to evaluate how enriched shared minor parent deserts and islands are relative to null expectations. One limitation of these permutations is that they do not precisely match the architecture of shared minor parent deserts and islands, and thus shared patterns of ancestry may be driven by residual effects of the local recombination and functional basepair environment (despite our attempts to control for this, see above).

As a third approach, for each minor parent island or desert identified in the Santa Cruz hybrid population, we calculated the length of the region in cMs and then summarized ancestry in windows of that size genome-wide. We next calculated the number or coding basepairs and the local recombination rate in each window. We identified all windows genome-wide that fell within 10% of the number of coding (or conserved basepairs) and within 10% of the local recombination rate of the focal minor parent ancestry island or desert. Next, we calculated ancestry for this subset of windows in the Santa Cruz population and in each *X. birchmanni* × *X. malinche* hybrid population. We performed 1,000 simulations randomly sampling a window from Santa Cruz and a window from each focal *X. birchmanni* × *X. malinche* hybrid population and asked how frequently shared minor parent ancestry islands or deserts were identified by chance. For minor parent ancestry deserts this was defined as falling in the lower 2.5% tail in Santa Cruz and lower 10% tail in *X. birchmanni* × *X. malinche* hybrid populations, as described in the main text.

The results of this analysis for each minor parent desert and island are shown in Table S14 & S15. We find that minor parent deserts and islands are unlikely to overlap with regions of low (or high) minor parent ancestry in other hybrid populations, consistent with the results of our permutations.

**Supporting Information 9. Power and evidence for other hybrid incompatibilities involving shared minor parent deserts**

In the main text we focus our scan for shared hybrid incompatibilities between *X. birchmanni* × *X. cortezi* and *X. birchmanni* × *X. malinche* species pairs on a minor parent ancestry desert that is shared across all replicate hybrid populations of both types. However, any shared ancestry desert across *X. birchmanni* × *X. cortezi* and *X. birchmanni* × *X. malinche* hybrid populations may be a candidate for a locus involved in shared hybrid incompatibilities across species.

To evaluate this possibility, we selected an ancestry informative marker covered in the *X. birchmanni* × *X. malinche* $F_2$ dataset from the center of each shared minor parent ancestry desert. We performed a scan against all other ancestry informative markers in the $F_2$ dataset (N=20,011) and calculated $\chi^2$ statistics for deviations from expected two-locus genotype frequencies (with four degrees of freedom). As described in the main text we used the observed genotype frequencies at each pair of markers to calculate expected two-locus genotype frequencies under independent segregation.

We explored two different approaches to determine significance thresholds for each ancestry desert. In the first approach, we permuted genotypes across individuals at a focal locus from the center of the empirically detected desert and then scanned for interactions in the rest of the genome. We repeated this permutation and genome-wide scan across 500 simulations to generate 500 null datasets. We took the lower 5 and 10% quantile of the minimum p-values from each simulation as our false positive rate threshold and asked whether any interactions detected in the real data exceeded the thresholds determined from simulations.

As a second approach, we wanted to account for the fact that genome-wide ancestry varies, even among $F_2$ hybrids which on average derive 50% of their genome from each parental species [25]. As a result, permuting the genotypes in simulations as described in the previous paragraph may bias us towards having a more permissive p-value threshold since we are breaking correlations in the data between any focal locus and that individual's genome-wide ancestry. To account for this, we used the genome-wide hybrid index for each $F_2$ to simulate genotypes at the focal locus using a random binomial weighted by that individual's admixture proportion. We then performed genome-wide scans across 500 replicate simulations and took the lower 5 and 10% quantile of the minimum p-values in each simulation, as described above. Indeed, we found that p-value thresholds were more conservative using this approach as opposed to a simple permutation-based approach. We thus proceeded with the more conservative thresholds in analysis of the real data.

In scanning for interactions using the empirical data, we found that the majority of shared deserts did not have interacting pairs detected in the *X. birchmanni* × *X. malinche* $F_2$ dataset that passed genome-wide multiple testing corrections, even at a relaxed threshold of a 10% false positive rate. The lowest p-value for interactions with each region investigated is listed in Table S16. However, we did identify two additional shared deserts with significant interactions detected at the 10% false positive rate threshold. One of these deserts appears to have a complex genetic architecture, showing evidence of interactions with two other regions of the genome.

We investigated possible functional links between regions with statistical evidence for interaction using STRING pathway analyses, as described in the main text (see Methods). We found that the complex interaction involving the shared desert on chromosome 5 harbored genes that are known to interact between with the partner regions (Fig. S24). By contrast, the interaction between the shared ancestry desert on chromosome 4 and chromosome 9 (Fig. 4C-D) did not harbor genes with known pathway interactions. However, there were several genes across the two regions that were co-expressed in our RNAseq dataset (see *Supporting Information 10. Gene network, co-expression, and gene ontology analysis of minor parent deserts and islands*), including the semaphorin genes SEMA6b and SEMA4b.

While we see evidence that 3 of the 21 shared ancestry deserts may be involved in hybrid incompatibilities (Fig. 4; Fig. S24), we do not find evidence for interactions involving the majority of shared deserts. This could indicate that other forms of selection act on these regions, or that we lack power to detect many of the interactions. To aid in the interpretation of these results, we investigated the strength of selection at which we have power to detect hybrid incompatibilities in our $F_2$ datasets using simulations. Using admix'em, we modeled classic Dobzhansky-Muller hybrid incompatibilities in 1,000 $F_2$ hybrids (sample size in our real data: N=943). As described for other admix'em simulations above, we modeled 24 chromosomes, 24,000 ancestry informative markers, and used the empirical recombination map for *X. birchmanni* to specify the recombination probabilities at sites along the chromosome. We simulated a range of selection coefficients ($s$=0.1, 0.5, 0.75, and 1) and modeled all hybrid incompatibilities as co-dominant ($h$=0.5). For each simulation, two selected sites involved in the incompatibility were randomly placed in the genome. We performed 100 replicate simulations for each selection coefficient. For each pair of selected sites, we ran $\chi^2$ tests as we had done for the real data (see above, Methods). To estimate power, we asked whether we detected the simulated hybrid incompatibility at a FPR of 10%.

We found that we had poor power to detect selection at moderate selection coefficients (6% at $s$=0.1) but that power increased with stronger selection on hybrids (13% at $s$=0.5, 85% at $s$=0.75). In the case of lethal hybrid incompatibilities, we estimated our power to detect these interactions was close to 100% (incompatibilities were detected in 100% of simulations with $s$=1). We note that we simulated a classic DMI [26] but that power may vary with different assumptions about the architecture and dominance of the hybrid incompatibility [27].

Given these results we conclude that we have limited power to detect DMIs under moderate selection using our $F_2$ data but good power to detect DMIs under extremely strong selection. This contrasts with the results of our power analysis for shared deserts, where we have good power to detect sites under more moderate selection (e.g. $s$=0.1; *Supporting Information 7.*

*Power to detect shared minor parent deserts*). This raises the possibility that a larger number of shared ancestry deserts are involved in hybrid incompatibilities but that we lack the power to detect them in our $F_2$ dataset. Exploring these regions in larger datasets or with an admixture mapping approach will be an exciting direction for future work.

**Supporting Information 10. Gene network, co-expression, and gene ontology analysis of minor parent deserts and islands**

In the main text we describe the identification of 21 shared minor parent deserts and 19 shared minor parent islands across *X. birchmanni* × *X. malinche* and *X. birchmanni* × *X. cortezi* hybrid populations. Because the size of the deserts and islands should be correlated with the strength of selection for or against minor parent ancestry in that region, it is difficult to meaningfully evaluate the genes that are most likely to drive the shared ancestry patterns of interest. Nonetheless, we report these genes in Table S17 & S18 and perform gene ontology and pathway enrichment analyses.

The average length of shared minor parent deserts in our dataset was 208 kb and the average length of minor parent islands was 249 kb. We lifted these coordinates over to the *X. maculatus* assembly using haltools [28] and identified ensembl gene ids for genes that fell in this interval. Next, we retrieved GO categories for each gene using the biomart package in R and performed a gene ontology test using GOstats with the entire genome as the gene universe and the minor parent deserts or islands as the focal gene set. This analysis yielded dozens of over-represented categories for both minor parent deserts and islands. Results of this analysis are reported in Table S19 & S20.

Since we identified at least one hybrid incompatibility that involves an interaction with one of the shared minor parent deserts, this suggests that other minor parent deserts are candidates for hybrid incompatibilities between species. A priori, we might expect genes involved in hybrid incompatibilities to be densely connected to other genes in their protein-protein interactions or in their expression networks [29,30].

Thus, we wanted to evaluate whether there was evidence that shared minor parent deserts (or islands) were more (or less) connected to other genes than expected by chance. To generate an appropriate null set for comparison, we took the lengths of minor parent deserts identified in the Santa Cruz hybrid population and randomly permuted these onto the genome to generate 1000 null datasets. We repeated this procedure for shared minor parent islands. Using these null datasets and the real data, we performed two sets of analyses.

We calculated total connectivity of genes in minor parent deserts and islands using the weighted co-expression analysis R package WGCNA [31] and compared the mean total connectivity of these genes to the distribution of mean total connectivity generated from the null simulations. Briefly, we used *kallisto* to pseudoalign reads from *X. birchmanni* and *X. malinche* brain tissue samples to a reference *X. birchmanni* transcriptome, and the R package DESeq2 [32] to normalize and apply a variance stabilizing transformation (VST) to gene counts. VST counts were used to calculate pairwise adjacency values between genes, using Pearson's correlation coefficient, an unsigned network model, and an optimal soft-threshold power of 7 (|cor|^7). The WGCNA function intramodularConnectivity was used to calculate the connectivity of each gene in relation to every other gene in the expression data set (total connectivity, or kTotal; the sum of adjacencies per gene). We found that the mean total connectivity of genes in both minor parent deserts (193) and islands (200) did not deviate from the null expectation; the 95% CI of mean total connectivity across the null datasets was 159 - 218 and 162 - 217, respectively (Fig. S34).

Finally, we assessed the connectivity of genes in minor parent deserts and islands using protein functional enrichment network data for *X. maculatus* from the STRING database [33]. Specifically, we obtained interaction counts from the experiments/biochemistry, co-expression, and combined score categories which had a confidence score above 0.4. Because direct experimental and co-expression evidence for *X. maculatus* is sparse in the STRING database, we used interaction counts from direct evidence and evidence for interaction discovered in other organisms and transferred through homology. We performed this step separately for the experiments and co-expression categories. The combined score category is a sum of all STRING-db categories and was not manipulated. An undirected graph was constructed from these network data by adding vertices corresponding to each Ensembl gene and edges corresponding to non-zero interaction counts between genes, using the networkx software [34]. Weighted degree centrality, a measure of the number of connections of each Ensembl gene weighted by the STRING-db interaction count corresponding to each interaction, was computed for each vertex (gene). Ensembl gene IDs were obtained for genes in the shared minor parent deserts and islands and the 1,000 matched permutation sets described previously. We then calculated the mean weighed degree centrality computed for all genes in the original and permutation sets (Figs. S35 and S36). We observed a tendency for lower connectivity in genes found in deserts (Fig. S35), although connectivity only significantly differed from the null background when considering the co-expression score category ($p$=0.038) and not the experiment-based ($p$=0.178) or the combined score ($p$=0.126) categories. We did not observe a significant difference in connectivity of genes in islands (Fig. S36) from the null background.

These findings run counter to our naïve expectation that highly connected genes might be enriched in shared deserts. This could suggest that apart from the mapped incompatibilities, other shared deserts and islands might not be explained by gene interactions. Alternately, interpretation of connectivity analysis may be limited by the paucity of direct protein interaction information in *X. maculatus,* since interaction scores were mostly inferred through homology. Finally, the assumption that hybrid incompatibilities are likely to have dense protein-protein interaction networks may itself be flawed, as it is a prediction based on first principles but difficult to evaluate in the absence of sufficient empirical data.

**Supporting Information 11. Analyzing correlations between recombination rate and ancestry after controlling for power to infer these values**

In the main text we present analysis of the relationship between the local recombination rate, inferred using a population genetic approach in *X. birchmanni*, and local ancestry in *X. birchmanni* × *X. cortezi* hybrids. However, our power to infer both recombination rate and ancestry varies along the genome with the density of informative sites. To ensure that the correlations between recombination rate and ancestry that we observe are not driven by these technical factors, we thinned our data before inferring recombination rate and ancestry as described in the main text, and repeated map estimates and HMM steps (see Methods).

Using the local ancestry data and recombination rates inferred from thinned input data, we next evaluated relationships between ancestry, recombination rate, and the number of linked functional basepairs in a partial correlation analysis at a range of physical non-overlapping window sizes (Table S2). Regardless of the exact analysis approach, our results were qualitatively similar (compare to Table S1). When we compared the results of our unthinned dataset to data that had been thinned by ancestry informative sites (AIMs) or analyzed with a thinned recombination rate map, we saw no substantive differences in the correlation between minor parent ancestry and recombination rate (Santa Cruz - 100 kb window unthinned: $\rho = 0.44$, $p<10^{-321}$, thinned AIMs: $\rho = 0.43$, $p<10^{-303}$, thinned recombination rate map: $\rho = 0.43$, $p<10^{-296}$, both thinned recombination rate map and thinned AIMs: $\rho = 0.41$, $p<10^{-278}$). We also explored other filtering approaches, such as excluding regions that might be impacted by adaptive introgression by removing windows that fell in the top 1% of minor parent ancestry. In doing so, we found that these high minor parent ancestry regions were not driving observed correlations between ancestry and recombination rate (Santa Cruz - 100 kb window: $\rho = 0.44$, $p<10^{-321}$ and removing the top 1%: $\rho = 0.44$, $p<10^{-323}$). To test if autocorrelation across the genome in ancestry and recombination rate affected our results we thinned our data by physical distance, retaining only one window per 500 kb. This also had no substantial effect on our findings (Santa Cruz - 100 kb window: $\rho = 0.44$, $p<10^{-321}$ and thinning to one window per 500 kb $\rho = 0.47$, $p<10^{-72}$). The observed correlations between minor parent ancestry and recombination rate did not vary substantially between Huextetitla and Santa Cruz (100 kb window Santa Cruz: $\rho = 0.44$, $p<10^{-321}$ and Huextetitla: $\rho = 0.42$, $p<10^{-283}$). See Table S2 for a summary of these results at different window sizes.

**Supporting Information 12. Shared minor parent deserts and islands using Huextetitla as the focal population**

In the main text, we identify shared minor parent islands and deserts between *X. birchmanni* × *X. cortezi* and *X. birchmanni* × *X. malinche* hybrid populations using Santa Cruz as our focal population. We made this choice because we had a much larger sample size for the Santa Cruz hybrid population (N=12 vs N=242), but results are qualitatively similar if we perform this analysis focusing instead on Huextetitla, as described below.

We identified minor parent deserts and islands in Huextetitla in the same manner as we did with Santa Cruz (see Methods). Since we have fewer individuals from the Huextetitla population, many more regions of the genome are inferred to be fixed for *X. cortezi* ancestry (major parent ancestry). This meant we only identified deserts among regions that are fixed for major parent ancestry in our sample, and that this number of regions is likely an inflation of the number of regions truly fixed for major parent ancestry in Huextetitla. We identified a total of 582 deserts; 109 of these were shared with Santa Cruz, and 28 of which were shared with one or more of the *X. birchmanni* × *X. malinche* hybrid populations. Permutations indicate that similar to the results presented in the main text, this degree of sharing of minor parent deserts is unexpected by chance. In permutations of our data we find that we expect fewer than 11 shared deserts by chance in pairwise comparisons across populations. Repeating this procedure for minor parent islands, we identified a total of 70 islands, 61 of which were shared in Santa Cruz, and 17 of which were shared in any three *X. birchmanni* × *X. malinche* hybrid populations. With permutations, we would expect less than seven to be shared by chance in pairwise comparisons across populations.

**Table S1.** Correlations between minor parent ancestry (*X. birchmanni* ancestry) and recombination rate in Santa Cruz and Huextetitla hybrid populations at different non-overlapping window sizes.

| **Population** | **Spearman's correlation between minor ancestry and recombination rate** | | |
|---|---|---|---|
| | **50 kb** | **100 kb** | **250 kb** |
| **Santa Cruz** | $\rho = 0.40$<br>$p < 10^{-325}$ | $\rho = 0.44$<br>$p = 10^{-321}$ | $\rho = 0.51$<br>$p = 10^{-180}$ |
| **Huextetitla** | $\rho = 0.37$<br>$p < 10^{-325}$ | $\rho = 0.42$<br>$p = 10^{-283}$ | $\rho = 0.50$<br>$p = 10^{-173}$ |

**Table S2.** Correlations between minor parent ancestry (*X. birchmanni* ancestry) and recombination rate in Santa Cruz and Huextetitla hybrid populations using different approaches to control for variation in power to infer local ancestry or to infer recombination rate. See Methods and Supporting Information 11 for more details on these analyses. AIMs – ancestry informative sites; Rec – recombination. In the mask short tracts analysis, we removed ancestry tracts shorter than 0.004 cM for minor parent ancestry tracts and 0.035 cM for major parent ancestry tracts, based on the reasoning that these unusually short tracts compared to expectations given the age of the hybrid population [35] might represent switch errors. In the thinned physical distance analysis, one window was retained every 500 kb. In the exclude inversions category, we removed chromosomes 21 and 24 which have large inversions between *X. birchmanni* and *X. cortezi*.

| **Population** | **Additional Analysis** | **Spearman's correlation between minor parent ancestry and recombination rate** | | |
|---|---|---|---|---|
| | | **50 kb** | **100 kb** | **250 kb** |
| **Santa Cruz** | thinned AIMs | $\rho = 0.39$<br>$p < 10^{-325}$ | $\rho = 0.43$<br>$p = 10^{-303}$ | $\rho = 0.49$<br>$p = 10^{-167}$ |
| | thinned Rec Map | $\rho = 0.38$<br>$p < 10^{-325}$ | $\rho = 0.43$<br>$p = 10^{-296}$ | $\rho = 0.50$<br>$p = 10^{-173}$ |
| | thinned AIMs & thinned Rec Map | $\rho = 0.38$<br>$p < 10^{-325}$ | $\rho = 0.41$<br>$p = 10^{-278}$ | $\rho = 0.49$<br>$p = 10^{-161}$ |
| | mask short tracts | $\rho = 0.40$<br>$p < 10^{-325}$ | $\rho = 0.44$<br>$p = 10^{-323}$ | $\rho = 0.51$<br>$p = 10^{-182}$ |
| | thinned physical distance | $\rho = 0.41$<br>$p = 10^{-56}$ | $\rho = 0.47$<br>$p = 10^{-72}$ | $\rho = 0.53$<br>$p = 10^{-96}$ |
| | exclude inversions | $\rho = 0.40$<br>$p < 10^{-325}$ | $\rho = 0.44$<br>$p = 10^{-301}$ | $\rho = 0.51$<br>$p = 10^{-170}$ |
| **Huextetitla** | thinned AIMs | $\rho = 0.36$<br>$p < 10^{-325}$ | $\rho = 0.40$<br>$p = 10^{-257}$ | $\rho = 0.58$<br>$p = 10^{-156}$ |
| | thinned Rec Map | $\rho = 0.35$<br>$p < 10^{-325}$ | $\rho = 0.40$<br>$p = 10^{-262}$ | $\rho = 0.49$<br>$p = 10^{-165}$ |
| | thinned AIMs & thinned Rec Map | $\rho = 0.34$<br>$p < 10^{-325}$ | $\rho = 0.38$<br>$p = 10^{-238}$ | $\rho = 0.47$<br>$p <= 10^{-149}$ |

**Table S3.** Correlations between minor parent ancestry (*X. birchmanni* ancestry) and recombination rate in Santa Cruz and Huextetitla hybrid populations using a $F_2$ crossover map to estimate recombination rate. See Methods for details.

| **Population** | **Additional Analysis** | **Spearman's correlation between minor parent ancestry and crossover rate per window** | |
|---|---|---|---|
| | | 1 Mb | 5 Mb |
| **Santa Cruz** | Recombination rate from observed crossovers in $F_2$s | $\rho = 0.64$<br>$p = 10^{-79}$ | $\rho = 0.69$<br>$p = 10^{-22}$ |
| **Huextetitla** | | $\rho = 0.63$<br>$p = 10^{-76}$ | $\rho = 0.64$<br>$p = 10^{-18}$ |

**Table S4.** Correlations between minor parent ancestry (*X. birchmanni* ancestry) and the number of coding and conserved basepairs in a range of genetic non-overlapping window sizes. 75% of ancestry tracts found in Santa Cruz and Huextetitla are 0.1 cM or larger.

| **Population** | **Spearman's partial correlation with minor parent ancestry** | | | | | | | |
|---|---|---|---|---|---|---|---|---|
| | **0.1 cM** | | **0.25 cM** | | **0.5 cM** | | **1 cM** | |
| | **Coding** | **Conserved** | **Coding** | **Conserved** | **Coding** | **Conserved** | **Coding** | **Conserved** |
| **Santa Cruz** | $\rho$=0.04 p = $10^{-8}$ | $\rho$= -0.17 p = $10^{-96}$ | $\rho$=0.06 p = $10^{-6}$ | $\rho$= -0.14 p= $10^{-54}$ | $\rho$=0.05 p= $10^{-3}$ | $\rho$= -0.18 p= $10^{-31}$ | $\rho$=0.05 p=0.025 | $\rho$= -0.22 p= $10^{-19}$ |
| **Huextetitla** | $\rho$=0.06 p =$10^{-11}$ | $\rho$= -0.14 p = $10^{-64}$ | $\rho$=0.07 p= $10^{-7}$ | $\rho$= -0.16 p= $10^{-37}$ | $\rho$=0.07 p= $10^{-4}$ | $\rho$= -0.22 p= $10^{-24}$ | $\rho$=0.06 p= $10^{-3}$ | $\rho$= -0.19 p= $10^{-15}$ |

**Table S5.** Analysis of the correlation between minor parent ancestry and linked coding and conserved basepairs in 0.25 cM non-overlapping windows, excluding all regions with structural rearrangements between *X. birchmanni*, *X. malinche*, or *X. cortezi*.

| Population | Spearman's correlation with minor parent ancestry | |
|---|---|---|
| | 0.25 cM | |
| | Coding | Conserved |
| Santa Cruz | $\rho$=-0.21<br>p = $10^{-59}$ | $\rho$= -0.28<br>p= $10^{-106}$ |
| Huextetitla | $\rho$=-0.15<br>p= $10^{-29}$ | $\rho$= -0.22<br>p= $10^{-61}$ |

**Table S6.** Relationship between minor parent ancestry (*X. birchmanni* ancestry) and the number of synonymous and nonsynonymous substitutions found in a non-overlapping window of a given genetic size. Ancestry was summarized using the results of an HMM run on a set of thinned input ancestry informative sites (see Methods). Minor parent ancestry is reduced in regions with a higher number of both synonymous and nonsynonymous substitutions between *X. birchmanni* and *X. malinche*. This may be driven by a correlation between coding substitutions and regions with a high number of linked coding basepairs (Table S7).

| **Population** | **nt change** | **Spearman's correlation with minor parent ancestry** | | | |
|---|---|---|---|---|---|
| | | **0.1 cM** | **0.25 cM** | **0.5 cM** | **1 cM** |
| **Santa Cruz** | non-synonymous | $\rho$ = -0.10<br>p = $10^{-30}$ | $\rho$ = -0.13<br>p = $10^{-23}$ | $\rho$ = -0.16<br>p = $10^{-20}$ | $\rho$ = -0.20<br>p = $10^{-16}$ |
| | synonymous | $\rho$ = -0.12<br>p = $10^{-49}$ | $\rho$ = -0.16<br>p = $10^{-36}$ | $\rho$ = -0.19<br>p = $10^{-28}$ | $\rho$ = -0.25<br>p = $10^{-24}$ |
| **Huextetitla** | non-synonymous | $\rho$ = -0.05<br>p = $10^{-11}$ | $\rho$ = -0.08<br>p = $10^{-10}$ | $\rho$ = -0.11<br>p = $10^{-10}$ | $\rho$ = -0.14<br>p = $10^{-9}$ |
| | synonymous | $\rho$ = -0.08<br>p = $10^{-21}$ | $\rho$ = -0.11<br>p = $10^{-17}$ | $\rho$ = -0.14<br>p = $10^{-15}$ | $\rho$ = -0.18<br>p = $10^{-14}$ |

**Table S7.** Relationship between minor parent ancestry (*X. birchmanni* ancestry), the number of coding basepairs, and the number of synonymous and nonsynonymous substitutions found in a window of a given genetic size. Ancestry was summarized using the results of an HMM run on a set of thinned input ancestry informative sites (see Methods).

| **Population** | **nt change** | **Spearman's partial correlation** | | | |
|---|---|---|---|---|---|
| | | **0.1 cM** | **0.25 cM** | **0.5 cM** | **1 cM** |
| **Santa Cruz** | non-synonymous | $\rho = 0.06$<br>$p = 10^{-12}$ | $\rho = 0.10$<br>$p = 10^{-15}$ | $\rho = 0.13$<br>$p = 10^{-13}$ | $\rho = 0.17$<br>$p = 10^{-12}$ |
| | synonymous | $\rho = 0.04$<br>$p = 10^{-7}$ | $\rho = 0.08$<br>$p = 10^{-11}$ | $\rho = 0.12$<br>$p = 10^{-11}$ | $\rho = 0.11$<br>$p = 10^{-6}$ |
| | coding | $\rho = -0.14$<br>$p = 10^{-64}$ | $\rho = -0.22$<br>$p = 10^{-66}$ | $\rho = -0.28$<br>$p = 10^{-60}$ | $\rho = -0.33$<br>$p = 10^{-44}$ |
| **Huextetitla** | non-synonymous | $\rho = 0.06$<br>$p = 10^{-11}$ | $\rho = 0.09$<br>$p = 10^{-12}$ | $\rho = 0.11$<br>$p = 10^{-10}$ | $\rho = 0.15$<br>$p = 10^{-9}$ |
| | synonymous | $\rho = 0.04$<br>$p = 10^{-05}$ | $\rho = 0.08$<br>$p = 10^{-10}$ | $\rho = 0.11$<br>$p = 10^{-10}$ | $\rho = 0.11$<br>$p = 10^{-6}$ |
| | coding | $\rho = -0.11$<br>$p = 10^{-40}$ | $\rho = -0.19$<br>$p = 10^{-49}$ | $\rho = -0.24$<br>$p = 10^{-43}$ | $\rho = -0.30$<br>$p = 10^{-35}$ |

**Table S8.** Analysis of the correlation between minor parent ancestry and linked coding and conserved basepairs in 0.25 cM non-overlapping windows. Ancestry was summarized using the results of an HMM run on a set of thinned input ancestry informative sites (see Methods).

| Population | Spearman's partial correlation with minor parent ancestry | |
|---|---|---|
| | 0.25 cM | |
| | Coding | Conserved |
| Santa Cruz | $\rho$ =0.05<br>p = $10^{-4}$ | $\rho$ = -0.20<br>p= $10^{-55}$ |
| Huextetitla | $\rho$ =0.06<br>p= $10^{-5}$ | $\rho$ = -0.17<br>p= $10^{-40}$ |

**Table S9.** Analysis of the correlation between minor parent ancestry and linked conserved basepairs in non-overlapping windows in a filtered dataset. Here, windows in the lower or upper 25% of ancestry informative sites were dropped to exclude windows where we had especially low or high power to infer ancestry.

| **Population** | **Spearman's partial correlation with minor parent ancestry** | | |
|---|---|---|---|
| | **0.05 cM** | **0.1 cM** | **0.25 cM** |
| **Santa Cruz** | $\rho = -0.1$ $p<10^{-36}$ | $\rho = -0.12$ $p<10^{-28}$ | $\rho = -0.17$ $p<10^{-22}$ |
| **Huextetitla** | $\rho = -0.07$ $p<10^{-18}$ | $\rho = -0.09$ $p<10^{-16}$ | $\rho = -0.13$ $p<10^{-14}$ |

**Table S10**. Cross-population correlations in minor parent ancestry at a range of non-overlapping window sizes, without controlling for recombination rate and coding/conserved basepair covariates. Santa Cruz and Huextetitla populations are *X. birchmanni* × *X. cortezi* hybrid populations that derive the majority of their genome from *X. cortezi*. Acuapa and Aguazarca are *X. birchmanni* × *X. malinche* hybrid populations that derive the majority of their genome from *X. birchmanni*; Tlatemaco is a *X. birchmanni* × *X. malinche* hybrid population that derives the majority of its genome from *X. malinche*.

| Population | Comparison Population | Spearman's correlation with minor parent ancestry | | |
|---|---|---|---|---|
| | | **50 kb** | **100 kb** | **250 kb** |
| **Santa Cruz** | Huextetitla | $\rho = 0.75$ <br> $p < 10^{-325}$ | $\rho = 0.78$ <br> $p < 10^{-325}$ | $\rho = 0.82$ <br> $p < 10^{-325}$ |
| **Santa Cruz** | Tlatemaco | $\rho = 0.01$ <br> $p = 0.15$ | $\rho = 0.02$ <br> $p = 0.19$ | $\rho = 0.03$ <br> $p = 0.15$ |
| **Santa Cruz** | Acuapa | $\rho = 0.23$ <br> $p = 10^{-159}$ | $\rho = 0.25$ <br> $p = 10^{-93}$ | $\rho = 0.28$ <br> $p = 10^{-49}$ |
| **Santa Cruz** | Aguazarca | $\rho = 0.15$ <br> $p = 10^{-74}$ | $\rho = 0.17$ <br> $p = 10^{-42}$ | $\rho = 0.19$ <br> $p = 10^{-23}$ |
| **Huextetitla** | Tlatemaco | $\rho = 0.02$ <br> $p = 0.06$ | $\rho = 0.02$ <br> $p = 0.14$ | $\rho = 0.03$ <br> $p = 0.12$ |
| **Huextetitla** | Acuapa | $\rho = 0.23$ <br> $p = 10^{-163}$ | $\rho = 0.25$ <br> $p = 10^{-97}$ | $\rho = 0.29$ <br> $p = 10^{-55}$ |
| **Huextetitla** | Aguazarca | $\rho = 0.15$ <br> $p = 10^{-75}$ | $\rho = 0.17$ <br> $p = 10^{-44}$ | $\rho = 0.19$ <br> $p= 10^{-24}$ |
| **Tlatemaco** | Acuapa | $\rho = -0.08$ <br> $p = 10^{-19}$ | $\rho = -0.07$ <br> $p = 10^{-9}$ | $\rho = -0.06$ <br> $p = 10^{-3}$ |
| **Tlatemaco** | Aguazarca | $\rho = -0.07$ <br> $p = 10^{-16}$ | $\rho = -0.07$ <br> $p = 10^{-8}$ | $\rho = -0.06$ <br> $p = 0.002$ |
| **Acuapa** | Aguazarca | $\rho = 0.36$ <br> $p < 10^{-325}$ | $\rho = 0.36$ <br> $p = 10^{-208}$ | $\rho = 0.38$ <br> $p = 10^{-92}$ |

**Table S11**. Cross-population correlations in minor parent ancestry at a range of non-overlapping window sizes, where windows containing rearrangements in any species have been removed, without controlling for recombination rate and coding/conserved basepair covariates. Santa Cruz and Huextetitla populations are *X. birchmanni* × *X. cortezi* hybrid populations that derive the majority of their genome from *X. cortezi*. Acuapa and Aguazarca are *X. birchmanni* × *X. malinche* hybrid populations that derive the majority of their genome from *X. birchmanni*; Tlatemaco is a *X. birchmanni* × *X. malinche* hybrid population that derives the majority of its genome from *X. malinche*.

| **Population** | **Comparison Population** | **Spearman's correlation with minor parent ancestry** | | |
|---|---|---|---|---|
| | | **50 kb** | **100 kb** | **250 kb** |
| **Santa Cruz** | Huextetitla | $\rho = 0.76$<br>$p < 10^{-325}$ | $\rho = 0.78$<br>$p < 10^{-325}$ | $\rho = 0.83$<br>$p < 10^{-325}$ |
| | Tlatemaco | $\rho = 0.00$<br>$p = 0.91$ | $\rho = 0.01$<br>$p = 0.67$ | $\rho = 0.02$<br>$p = 0.35$ |
| | Acuapa | $\rho = 0.23$<br>$p = 10^{-150}$ | $\rho = 0.24$<br>$p = 10^{-87}$ | $\rho = 0.28$<br>$p = 10^{-46}$ |
| | Aguazarca | $\rho = 0.16$<br>$p = 10^{-71}$ | $\rho = 0.17$<br>$p = 10^{-41}$ | $\rho = 0.19$<br>$p = 10^{-22}$ |
| **Huextetitla** | Tlatemaco | $\rho = 0.01$<br>$p = 0.14$ | $\rho = 0.02$<br>$p = 0.23$ | $\rho = 0.03$<br>$p = 0.16$ |
| | Acuapa | $\rho = 0.23$<br>$p = 10^{-153}$ | $\rho = 0.25$<br>$p = 10^{-90}$ | $\rho = 0.29$<br>$p = 10^{-51}$ |
| | Aguazarca | $\rho = 0.15$<br>$p = 10^{-68}$ | $\rho = 0.17$<br>$p = 10^{-40}$ | $\rho = 0.19$<br>$p= 10^{-22}$ |
| **Tlatemaco** | Acuapa | $\rho = -0.09$<br>$p = 10^{-26}$ | $\rho = -0.09$<br>$p = 10^{-13}$ | $\rho = -0.09$<br>$p = 10^{-6}$ |
| | Aguazarca | $\rho = -0.08$<br>$p = 10^{-19}$ | $\rho = -0.08$<br>$p = 10^{-10}$ | $\rho = -0.08$<br>$p = 10^{-5}$ |
| **Acuapa** | Aguazarca | $\rho = 0.35$<br>$p < 10^{-325}$ | $\rho = 0.35$<br>$p = 10^{-185}$ | $\rho = 0.36$<br>$p = 10^{-80}$ |

**Table S12.** Cross-population correlations in minor parent ancestry at a range of non-overlapping window sizes, including recombination rate, coding, and conserved basepair covariates. For each population and window size comparison, Spearman's $\rho$ from cross-population ancestry correlations after accounting for other features is given to the left of each cell. The estimated $\rho$ from the same partial correlation analysis of other features (recombination rate, number of coding basepairs, and number of conserved basepairs) are given one the right.

| **Population** | **Cross Population** | **Spearman's partial correlation with minor parent ancestry** | | | | | |
|---|---|---|---|---|---|---|---|
| | | **50 kb** | | **100 kb** | | **250 kb** | |
| **Santa cruz** | Huextetitla | $\rho = 0.71$<br>$p<10^{-325}$ | rec $\rho = 0.20$<br>coding $\rho = 0.04$<br>conserved $\rho = -0.03$ | $\rho = 0.73$<br>$p<10^{-325}$ | rec $\rho = 0.20$<br>coding $\rho = 0.04$<br>conserved $\rho = -0.03$ | $\rho = 0.77$<br>$p<10^{-325}$ | rec $\rho = 0.19$<br>coding $\rho = 0.04$<br>conserved $\rho = -0.03$ |
| | Tlatemaco | $\rho = 0.00$<br>$p = 0.88$ | rec $\rho = 0.39$<br>coding $\rho = 0.10$<br>conserved $\rho = -0.08$ | $\rho = 0.00$<br>$p = 0.85$ | rec $\rho = 0.43$<br>coding $\rho = 0.11$<br>conserved $\rho = -0.09$ | $\rho = 0.01$<br>$p = 0.63$ | rec $\rho = 0.50$<br>coding $\rho = 0.11$<br>conserved $\rho = -0.10$ |
| | Acuapa | $\rho = 0.15$<br>$p = 10^{-64}$ | rec $\rho = 0.36$<br>coding $\rho = 0.10$<br>conserved $\rho = -0.08$ | $\rho = 0.15$<br>$p = 10^{-33}$ | rec $\rho = 0.39$<br>coding $\rho = 0.11$<br>conserved $\rho = -0.09$ | $\rho = 0.16$<br>$p = 10^{-16}$ | rec $\rho = 0.46$<br>coding $\rho = 0.11$<br>conserved $\rho = -0.10$ |
| | Aguazarca | $\rho = 0.10$<br>$p = 10^{-30}$ | rec $\rho = 0.38$<br>coding $\rho = 0.10$<br>conserved $\rho = -0.08$ | $\rho = 0.09$<br>$p = 10^{-15}$ | rec $\rho = 0.42$<br>coding $\rho = 0.10$<br>conserved $\rho = -0.09$ | $\rho = 0.10$<br>$p = 10^{-7}$ | rec $\rho = 0.49$<br>coding $\rho = 0.11$<br>conserved $\rho = -0.09$ |
| **Huextetitla** | Santa Cruz | $\rho = 0.71$<br>$p<10^{-325}$ | rec $\rho = 0.11$<br>coding $\rho = 0.05$<br>conserved $\rho = -0.04$ | $\rho = 0.73$<br>$p<10^{-325}$ | rec $\rho = 0.13$<br>coding $\rho = 0.05$<br>conserved $\rho = -0.05$ | $\rho = 0.77$<br>$p <10^{-325}$ | rec $\rho = 0.16$<br>coding $\rho = 0.05$<br>conserved $\rho = -0.05$ |
| | Tlatemaco | $\rho = 0.01$<br>$p = 0.48$ | rec $\rho = 0.36$<br>coding $\rho = 0.11$<br>conserved $\rho = -0.09$ | $\rho = 0.01$<br>$p = 0.64$ | rec $\rho = 0.41$<br>coding $\rho = 0.11$<br>conserved $\rho = -0.10$ | $\rho = 0.01$<br>$p = 0.51$ | rec $\rho = 0.49$<br>coding $\rho = 0.12$<br>conserved $\rho = -0.10$ |
| | Acuapa | $\rho = 0.16$<br>$p = 10^{-73}$ | rec $\rho = 0.32$<br>coding $\rho = 0.11$<br>conserved $\rho = -0.09$ | $\rho = 0.16$<br>$p = 10^{-39}$ | rec $\rho = 0.37$<br>coding $\rho = 0.12$<br>conserved $\rho = -0.10$ | $\rho = 0.18$<br>$p = 10^{-21}$ | rec $\rho = 0.45$<br>coding $\rho = 0.12$<br>conserved $\rho = -0.10$ |
| | Aguazarca | $\rho = 0.10$<br>$p = 10^{-33}$ | rec $\rho = 0.35$<br>coding $\rho = 0.10$<br>conserved $\rho = -0.08$ | $\rho = 0.10$<br>$p = 10^{-17}$ | rec $\rho = 0.39$<br>coding $\rho = 0.11$<br>conserved $\rho = -0.09$ | $\rho = 0.10$<br>$p=10^{-7}$ | rec $\rho = 0.48$<br>coding $\rho = 0.11$<br>conserved $\rho = -0.10$ |

**Table S13.** Cross-population correlations in minor parent ancestry at a range of genetic non-overlapping window sizes.

| Population | Cross Population | Spearman's correlation with minor parent ancestry | | | |
|---|---|---|---|---|---|
| | | **0.1 cM** | **0.25 cM** | **0.5 cM** | **1 cM** |
| **Santa Cruz** | Huextetitla | $\rho = 0.87$<br>$p < 10^{-325}$ | $\rho = 0.89$<br>$p < 10^{-325}$ | $\rho = 0.91$<br>$p < 10^{-325}$ | $\rho = 0.93$<br>$p < 10^{-325}$ |
| | Tlatemaco | $\rho = -0.03$<br>$p = 10^{-5}$ | $\rho = -0.03$<br>$p = 0.03$ | $\rho = -0.01$<br>$p = 0.49$ | $\rho = 0.00$<br>$p = 0.94$ |
| | Acuapa | $\rho = 0.26$<br>$p = 10^{-205}$ | $\rho = 0.28$<br>$p = 10^{-106}$ | $\rho = 0.30$<br>$p = 10^{-66}$ | $\rho = 0.33$<br>$p = 10^{-43}$ |
| | Aguazarca | $\rho = 0.19$<br>$p = 10^{-105}$ | $\rho = 0.20$<br>$p = 10^{-57}$ | $\rho = 0.23$<br>$p = 10^{-37}$ | $\rho = 0.26$<br>$p = 10^{-27}$ |
| **Huextetitla** | Santa Cruz | $\rho = 0.87$<br>$p < 10^{-325}$ | $\rho = 0.89$<br>$p < 10^{-325}$ | $\rho = 0.91$<br>$p < 10^{-325}$ | $\rho = 0.93$<br>$p < 10^{-325}$ |
| | Tlatemaco | $\rho = -0.03$<br>$p = 0.003$ | $\rho = -0.01$<br>$p = 0.30$ | $\rho = 0.1$<br>$p = 0.69$ | $\rho = 0.02$<br>$p = 0.54$ |
| | Acuapa | $\rho = 0.22$<br>$p = 10^{-152}$ | $\rho = 0.24$<br>$p = 10^{-81}$ | $\rho = 0.27$<br>$p = 10^{-52}$ | $\rho = 0.30$<br>$p = 10^{-36}$ |
| | Aguazarca | $\rho = 0.17$<br>$p = 10^{-84}$ | $\rho = 0.19$<br>$p = 10^{-49}$ | $\rho = 0.22$<br>$p= 10^{-35}$ | $\rho = 0.25$<br>$p= 10^{-25}$ |

**Table S14 – S15 are provided as supplementary files**

**Table S16.** Strongest observed deviations from $\chi^2$ expectations in scans for interactors with each shared ancestry desert in $F_2$ data derived from *X. birchmanni* × *X. malinche* hybrids. Only deserts with interaction p-values $<10^{-3}$ are listed.

| Desert chromosome | Desert interval | Peak interacting chromosome: marker | Minimum $\chi^2$ p-value |
|---|---|---|---|
| ScyDAA6-1107-HRSCAF-1306 | 15626581-16090938 | ScyDAA6-932-HRSCAF-1100 7190108 | $7.3 \times 10^{-5}$ |
| ScyDAA6-1854-HRSCAF-2213 | 12052806-12133103 | ScyDAA6-8-HRSCAF-51 6886194 | $5.3 \times 10^{-5}$ |
| ScyDAA6-2188-HRSCAF-2635 | 22106437-22362200 | ScyDAA6-8-HRSCAF-51 18145420 | $1.2 \times 10^{-4}$ |
| ScyDAA6-2-HRSCAF-26 | 21866739-21867081 | ScyDAA6-2113-HRSCAF-2539 7620035 | $2.8 \times 10^{-4}$ |
| ScyDAA6-5984-HRSCAF-6694 | 10814724-10965078 | ScyDAA6-1934-HRSCAF-2318 15330451 | $2.1 \times 10^{-4}$ |
| ScyDAA6-7-HRSCAF-50 | 3461-24281 | ScyDAA6-1196-HRSCAF-1406 16551294 | $5.1 \times 10^{-5}$ |

**Table S17 – S20 are provided as supplementary files**

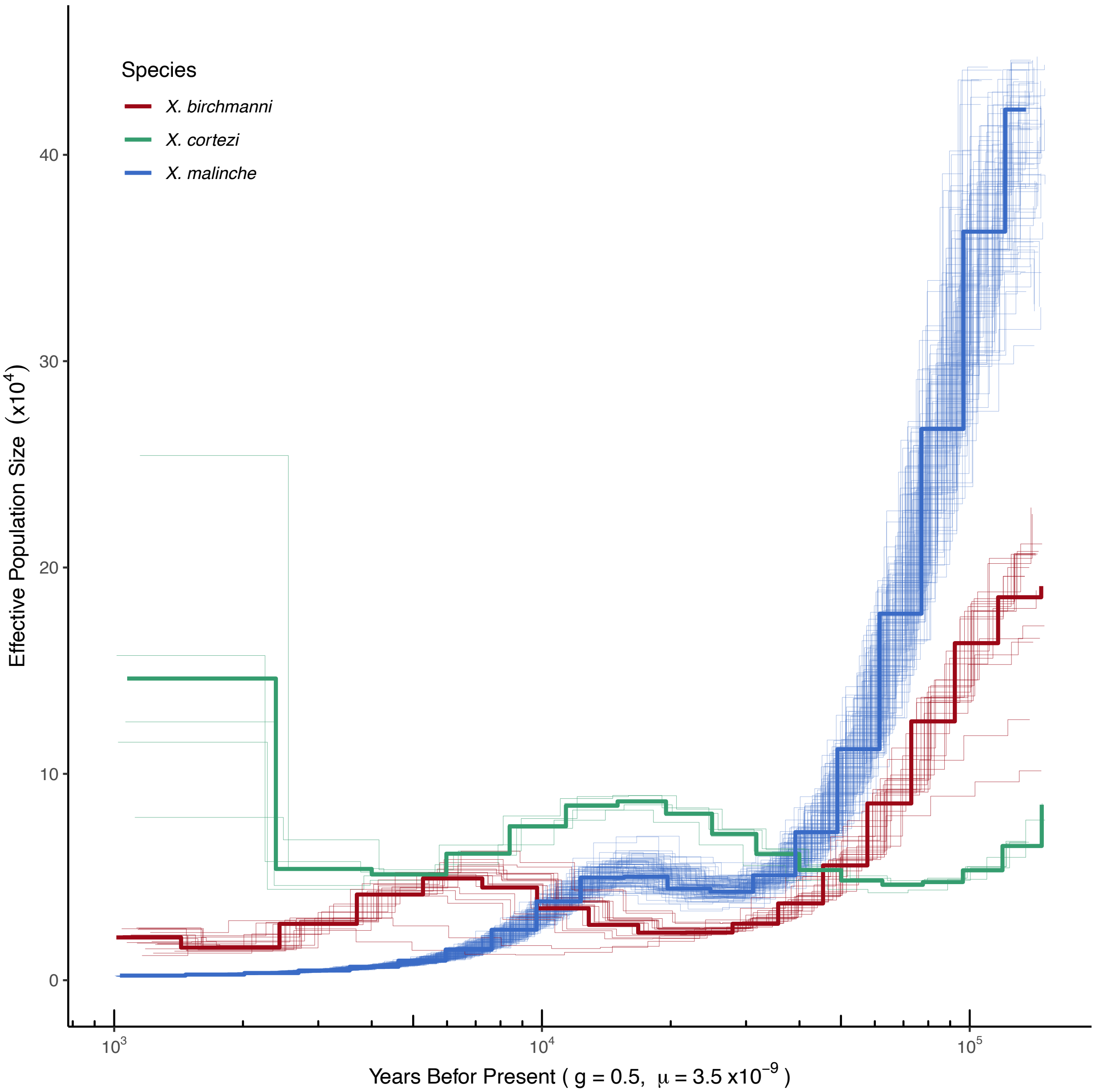


**Fig. S1.** PSMC results analyzing population history from a single whole-genome sample of *X. malinche* from the sampling site Chicayotla, 18 *X. birchmanni* individuals from the sampling site Coacuilco, and five *X. cortezi* individuals from the sampling site El Nacemiento de Huichihuayán. For visualization the single *X. malinche* sample was bootstrapped 100 times by resampling with replacement from the genome split into 500 kb segments. Analysis was conducted similarly to [5] with the time segmentation parameter set to 4+25*2+4+6, a $\rho/\theta$ ratio of 2, generation time of two generations per year, and mutation rate of $3.5 \times 10^{-9}$.

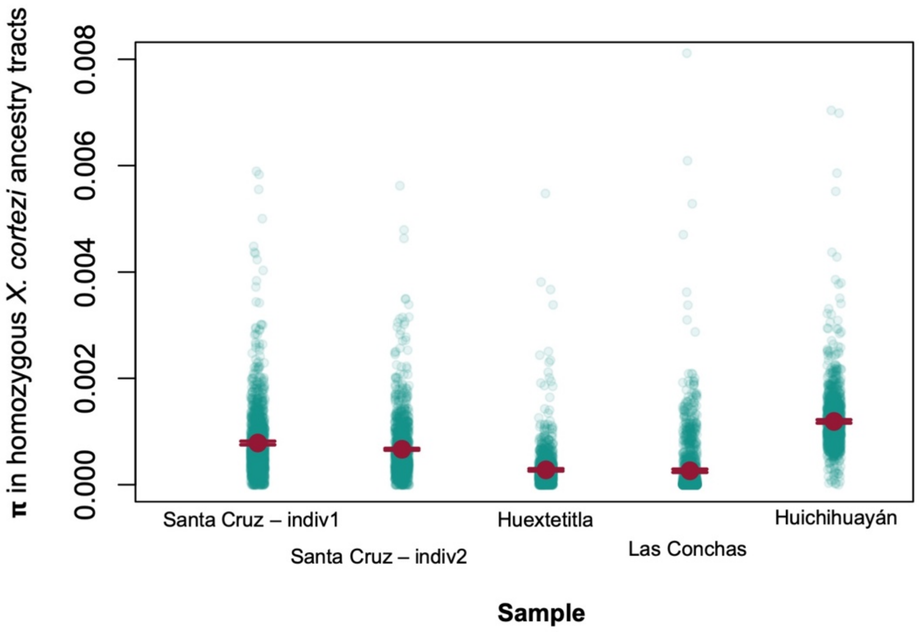


**Fig. S2.** Genetic diversity in homozygous *X. cortezi* ancestry tracts in individuals from two hybrid populations (Santa Cruz and Huextetitla) and in individuals from nearby allopatric *X. cortezi* parental populations (Las Conchas and Huichihuayán).

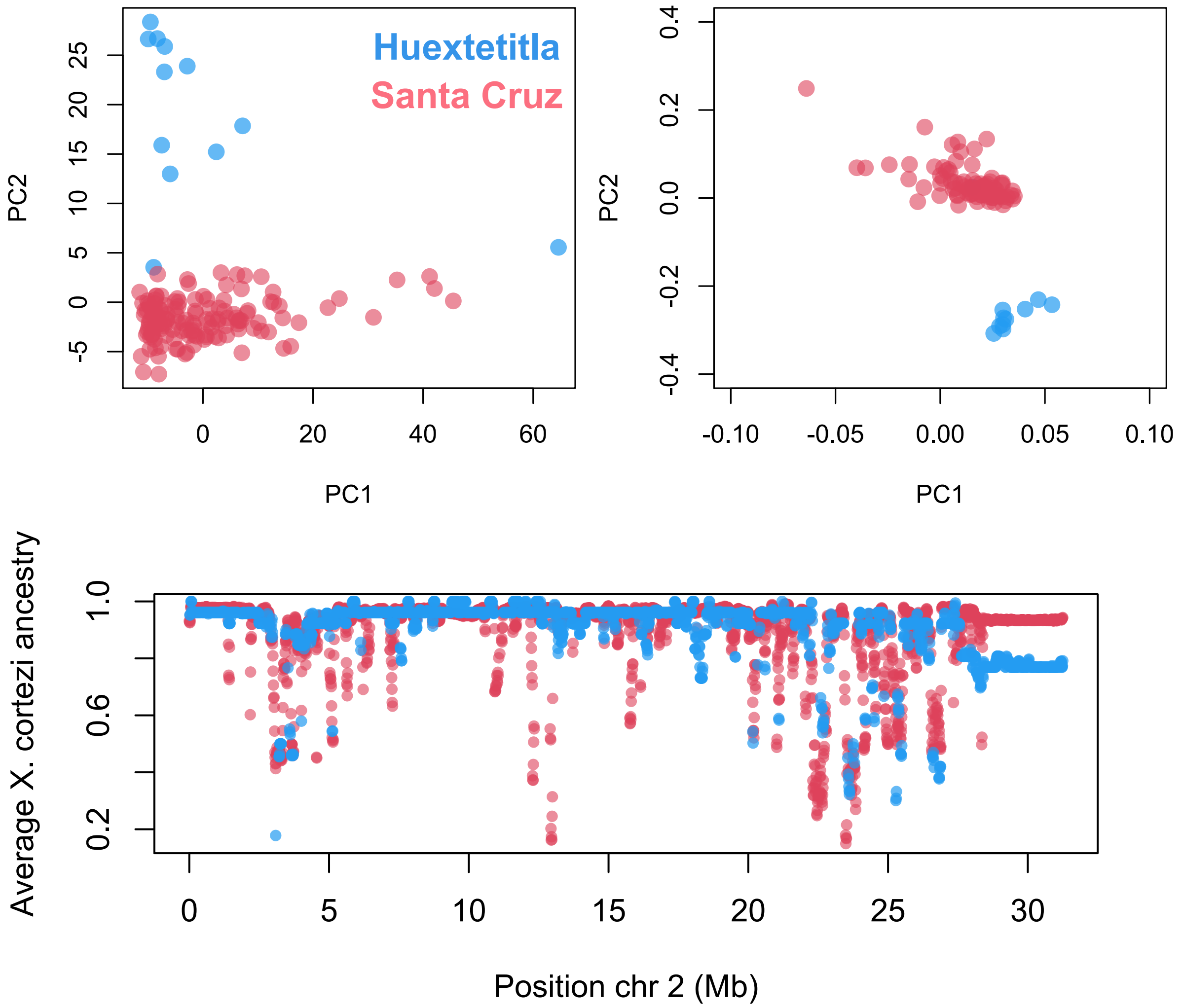


**Fig. S3.** (**Top**) (Left) PCA analysis of the locations of observed ancestry transitions in individuals from the Santa Cruz and Huextetitla hybrid populations. (Right) PCA analysis of pseudohaploid SNP calls derived from low-coverage sequence data of individuals from the Santa Cruz and Huextetitla populations (see Supporting Information 2). Separation along PC2 suggests that the Santa Cruz and Huextetitla hybrid populations have been somewhat independent in their recent demographic histories. (**Bottom**) Example of heterogeneity in ancestry along chromosome 2 in Huextetitla (blue) and Santa Cruz (pink) *X. birchmanni* × *X. cortezi* hybrid populations, averaged in 10 kb windows). Both populations have regions that are fixed or nearly fixed for both *X. cortezi* and *X. birchmanni* ancestry. Note the strong correlations in local ancestry between the two populations ($\rho$=0.65, p<$10^{-100}$).

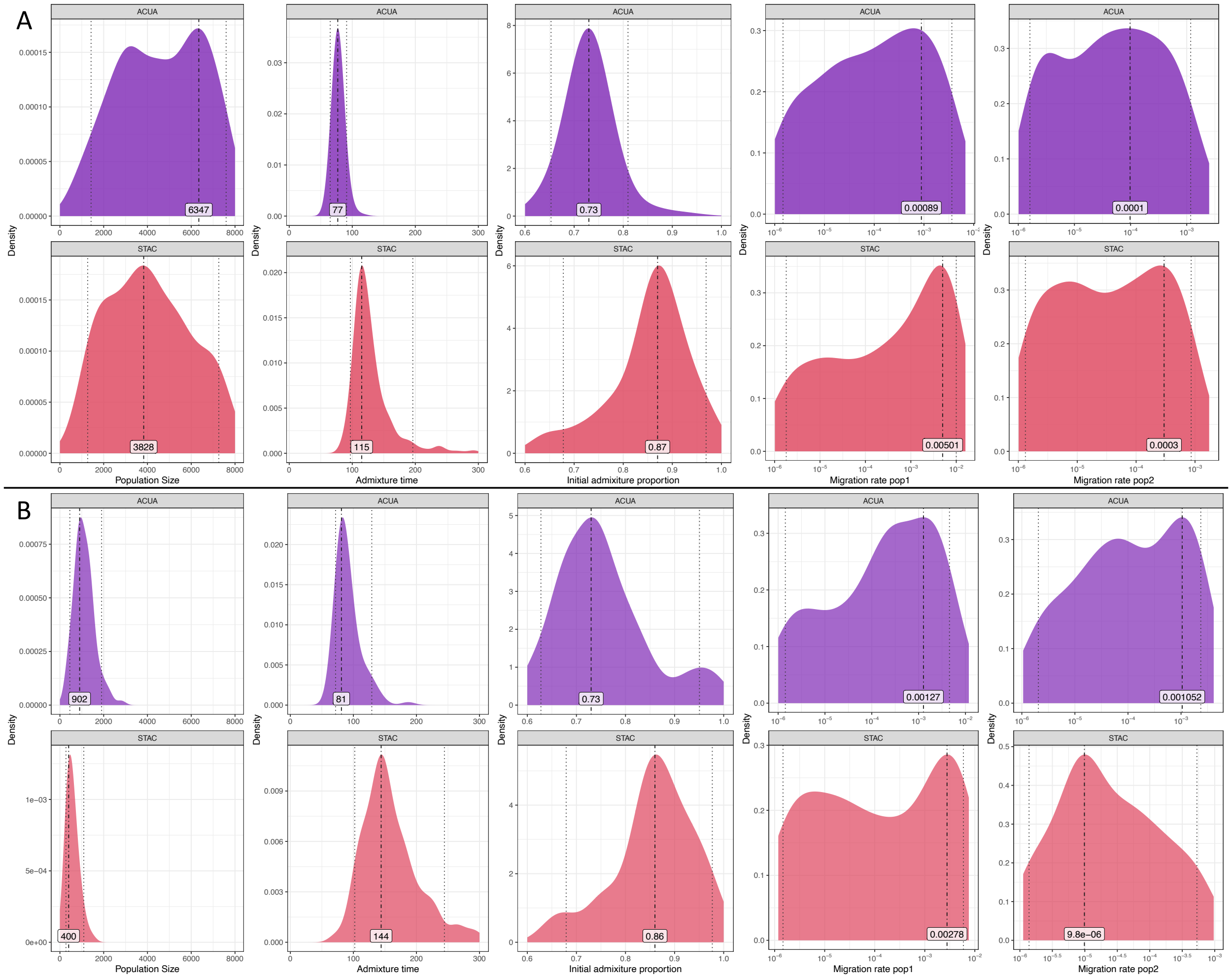


**Fig. S4.** Posterior distributions from Approximate Bayesian Computation (ABC) simulations used to infer the demographic history for a *X. birchmanni* × *X. cortezi* hybrid population Santa Cruz (STAC) (red) and a *X. birchmanni* × *X. malinche* hybrid population Acuapa (ACUA) (pruple). Dot-dashed lines and listed values correspond to the maximum a posteriori or MAP estimate for each distribution. Dotted lines are the 95% quantile range. See Supporting Information 3 for complete details of SLiM simulations and rejection sampling approach. Posterior distributions shown here are derived from uniform (or log-uniform) prior distributions of: initial population size, time since admixture (in generations), initial admixture proportion, and migration rate from each parent species. We accepted simulations based on two sets of summary statistics. **A.** Our primary analysis included summary statistics for the median length of minor parent ancestry tracts, average hybrid index, and the coefficient of variation for chromosome-wide ancestry across sampled individuals (ACUA N=500, STAC N=500). **B.** In a second analysis we included summary statistics for the median length of minor parent ancestry tracts, average hybrid index, and the coefficient of variation for local ancestry along the chromosome in 250 kb windows. We accepted very few simulations using the second approach (ACUA N=98, STAC N=90). We show the accepted simulations here to emphasize that the posterior distributions for most parameters are similar using the two approaches but rely on the inferences made in **A** for almost all analyses. See Supporting Information 3 for additional information.

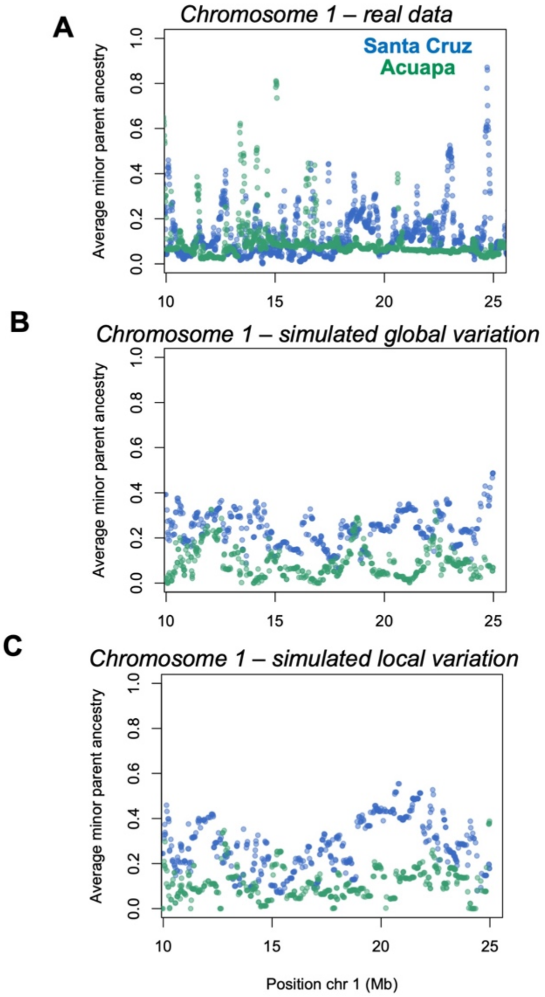


**Fig. S5**. **A.** Heterogeneity in minor parent ancestry in the real data for a section of chromosome 1 in the Santa Cruz and Acuapa populations. Points show the average minor parent ancestry in 10 kb windows. **B.** Results of one replicate simulation of local ancestry on chromosome 1 for Santa Cruz and Acuapa based on randomly drawn set of demographic parameters from the posterior distribution of ABC simulations that used global variation in admixture proportion as a summary statistic (see Supporting Information 3 & Fig. S4 for details). **C.** Results of one replicate simulation of local ancestry on chromosome 1 for Santa Cruz and Acuapa based on randomly drawn set of demographic parameters from the posterior distribution of ABC simulations that used local variation in admixture proportion (summarized in 250 kb windows) as a summary statistic. Points show the average minor parent ancestry in 10 kb windows. While these simulations incorporated inferred demographic history for each population they did not implement selection. This results in lower heterogenetity in local ancestry compared to the real data, even when a summary statistic of local variation in admixture proportion was used to accept or reject simulations (see Supporting Information 3 & Fig. S4).

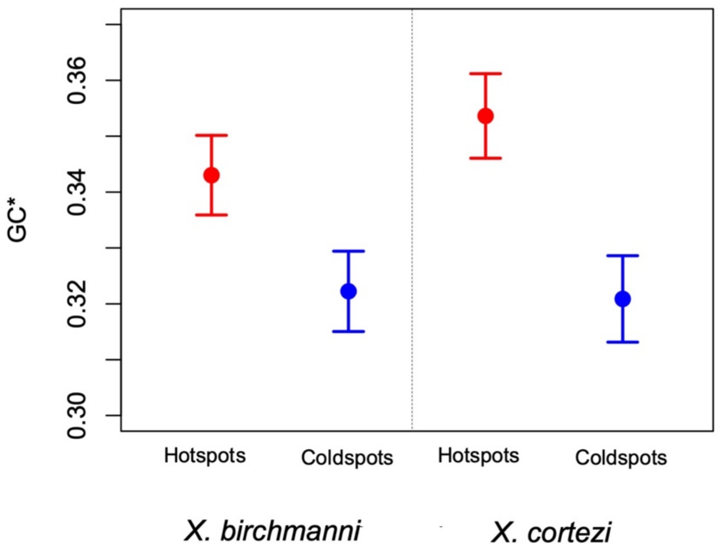


**Fig. S6.** GC*, a measure of GC biased gene conversion, in 5 kb hotspots identified in *X. birchmanni* as well has GC-content matched coldspots. Also shown is GC* for *X. cortezi* in the same hotspots and matched coldspots identified in *X. birchmanni*. These patterns suggest an excess of GC-biased gene conversion in hotspots identified in *X. birchmanni* in both species, providing further evidence that the fine scale recombination maps are shared between species.

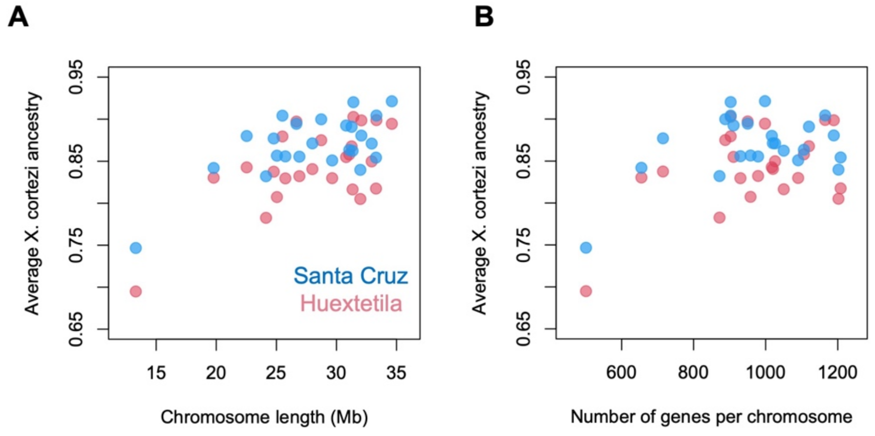


**Fig. S7.** In addition to heterogeneity in ancestry within chromosomes, we also observe substantial heterogeneity in ancestry proportion between chromosomes. **A.** In both Santa Cruz and Huextetitla this heterogeneity is modestly correlated with chromosome length, suggesting that it may be driven by higher effective recombination rates on shorter chromosomes ($\rho_{\text{Santa Cruz}} = 0.43$, $p_{\text{Santa Cruz}} = 0.036$; $\rho_{\text{Huextetitla}} = 0.39$, $p_{\text{Huextetitla}} = 0.058$). **B.** The correlation between number of genes per chromosome and chromosome-level ancestry is substantially weaker ($\rho_{\text{Santa Cruz}} = 0.12$, $p_{\text{Santa Cruz}} = 0.58$; $\rho_{\text{Huextetitla}} = 0.03$, $p_{\text{Huextetitla}} = 0.90$).

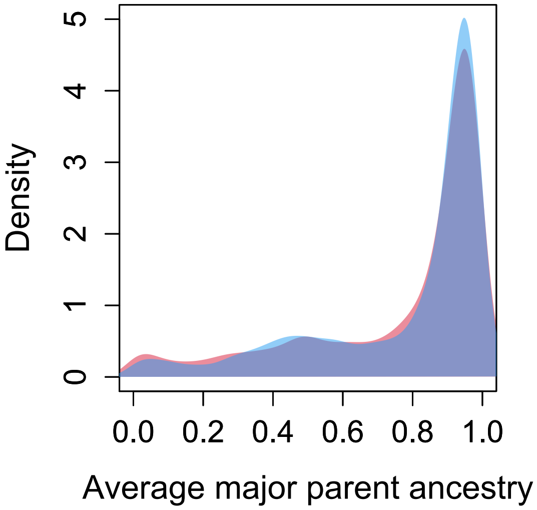


**Fig. S8.** No evidence for unusual minor parent ancestry in the Santa Cruz hybrid population in 0.1 cM windows with high nonsynonymous substitution rates between *X. cortezi* and *X. birchmanni* (upper 25% genome wide), versus matched windows with no nonsynonymous substitutions but a similar overall coding substitution rate. For each 0.1 cM window with a high number of nonsynonymous substitutions, we identified a 0.1 cM window with no nonsynonymous substitutions but an overall coding substitution rate (i.e. of synonymous substitutions) within 80-120% of that observed in the focal window. We see no significant differences in the minor parent ancestry distributions of the focal (pink) and matched (blue) windows.

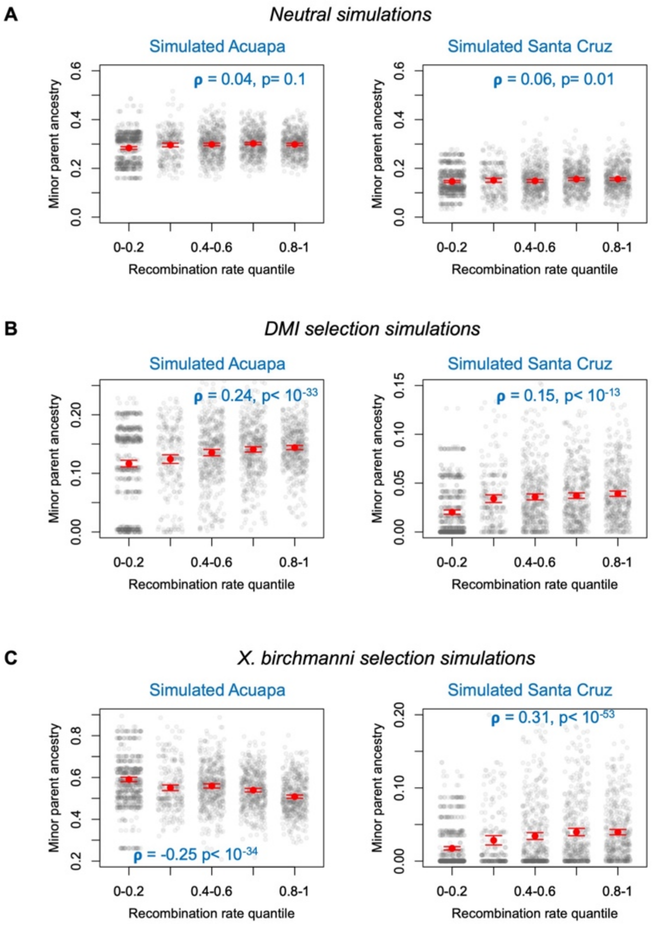


**Fig. S9.** Different models of selection on hybrids generate distinct correlations between minor parent ancestry and recombination rate. **A.** In the absence of selection there is no expected relationship between recombination rate and ancestry, and indeed this is what is observed in simulated Santa Cruz and Acuapa populations (single simulation example shown here). Gray points show minor parent ancestry in 250 kb windows, red points and whiskers show the mean and two standard errors of the mean. Inset shows correlation coefficient and p-value for the representative simulation. **B.** In the presence of selection against hybrid incompatibilities, selection drives a positive correlation between minor parent ancestry and recombination rate, regardless of the identity of the major parent species. Shown here are single representative simulations modeling the demographic history of the Santa Cruz and Acuapa populations with incompatibility selection implemented at 20 random pairs of sites throughout the genome. **C**. In the presence of selection against one parent species or the other, we expect to see conflicting directions in the correlation between minor parent ancestry and recombination rate depending on the admixture proportion of the hybrid population. In this set of simulations, a subset of sites derived from the *X. birchmanni* parent were globally disadvantageous, driving different patterns in the majority *X. birchmanni* (Acuapa) and minority *X. birchmanni* (Santa Cruz) hybrid populations. Simulations are described in detail in Supporting Information 6.

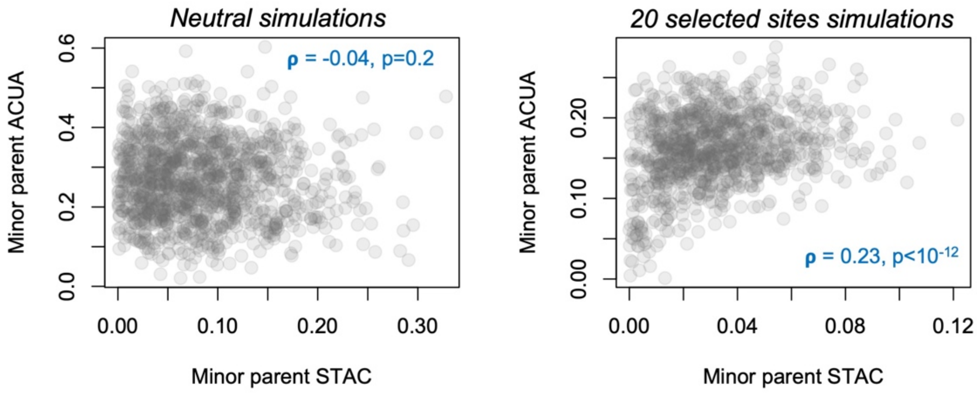


**Fig. S10.** Simulations suggest that we do not expect to observe cross-population correlations in local ancestry in the absence of shared sites under selection. **A.** Example simulation of local ancestry in Santa Cruz and Acuapa populations modeling inferred demographic history but no selection. Inset shows correlation coefficient and p-value for the pair of simulations. **B.** Example simulation of local ancestry in Santa Cruz and Acuapa populations modeling inferred demographic history and 20 randomly placed shared sites under selection in the two populations. Blue text shows correlation coefficient and p-value for the pair of simulations.

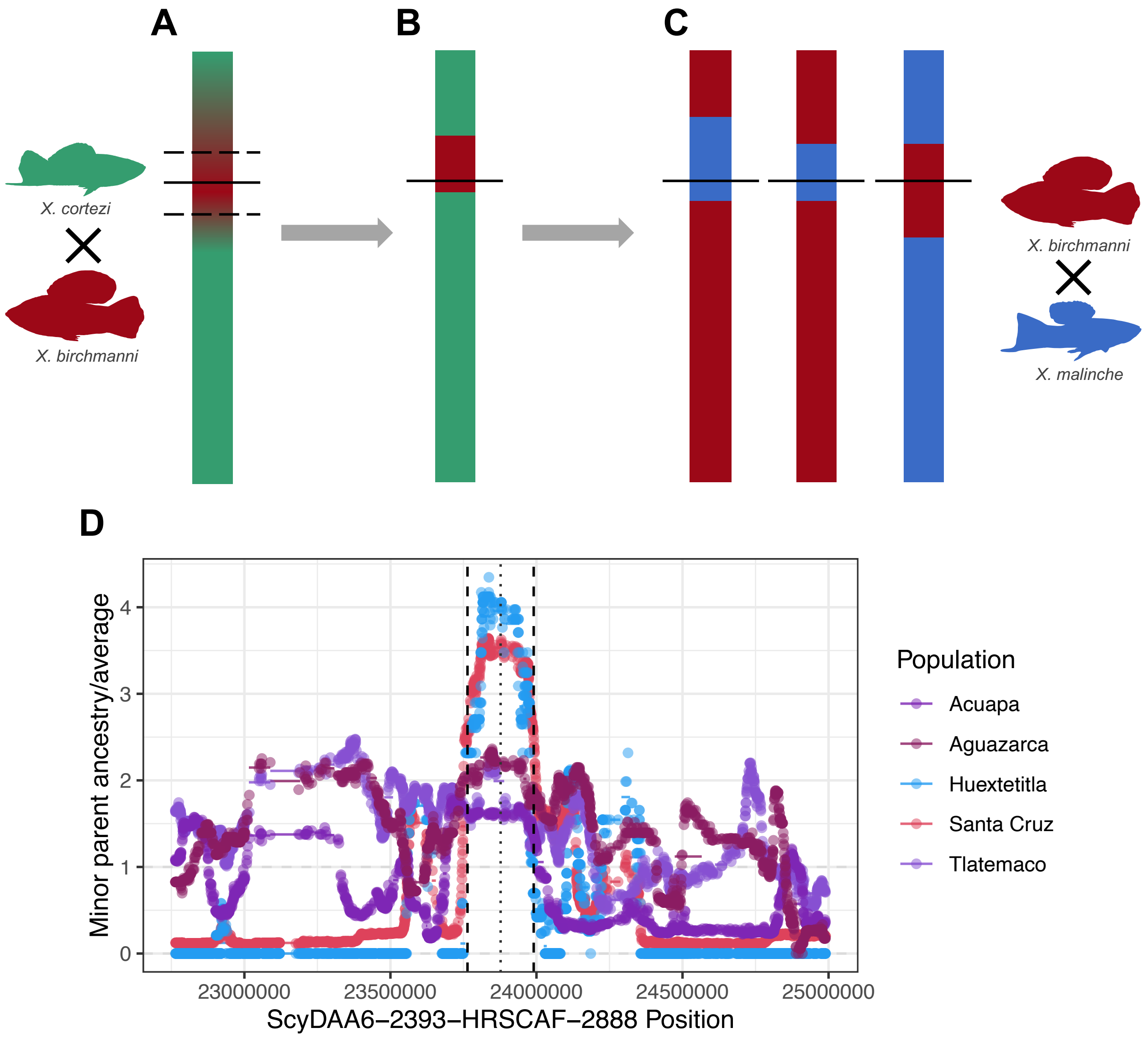


**Fig. S11.** Schematic of approach used to identify shared minor parent islands and deserts. We employed a stepwise approach to identify deserts and islands of minor parent ancestry and determine if they were shared across populations. Shown here is a hypothetical workflow for identifying a shared minor parent ancestry island. **A.** We started with identifying AIMs where average minor parent ancestry at that site exceeded the 97.5% quantile of minor parent ancestry genome-wide. From that focal site we expanded outward in the 5' and 3' directions to identify where minor parent ancestry falls below the 95% tail of the genome-wide distribution. This set the boundary of the focal minor parent island region. We then determined the midpoint of each region and identified the 0.05 cM window that contains the midpoint. **B.** We checked that the focal population's minor parent ancestry is greater than the 90% quantile of minor parent ancestry genome-wide when averaged across this 0.05 cM window. We then asked if this region is a shared minor parent ancestry outlier in other populations. **C.** Specifically, we evaluated minor parent ancestry in the midpoint 0.05 cM window in other hybrid populations. If minor parent ancestry in these populations is exceeded the 90% quantile of that population's genome wide ancestry distribution we classified that region as a shared minor parent island. **D.** Example of a minor parent island detected with this work flow. Dashed lines are the identified boundaries of the island and dotted line is the midpoint. Colored dots correspond to the minor parent ancestry at a given ancestry informative site divided by the genome wide average for that population. Colored lines indicate the minor parent ancestry for the focal 0.05 cM window divided by the genome wide average for the population.

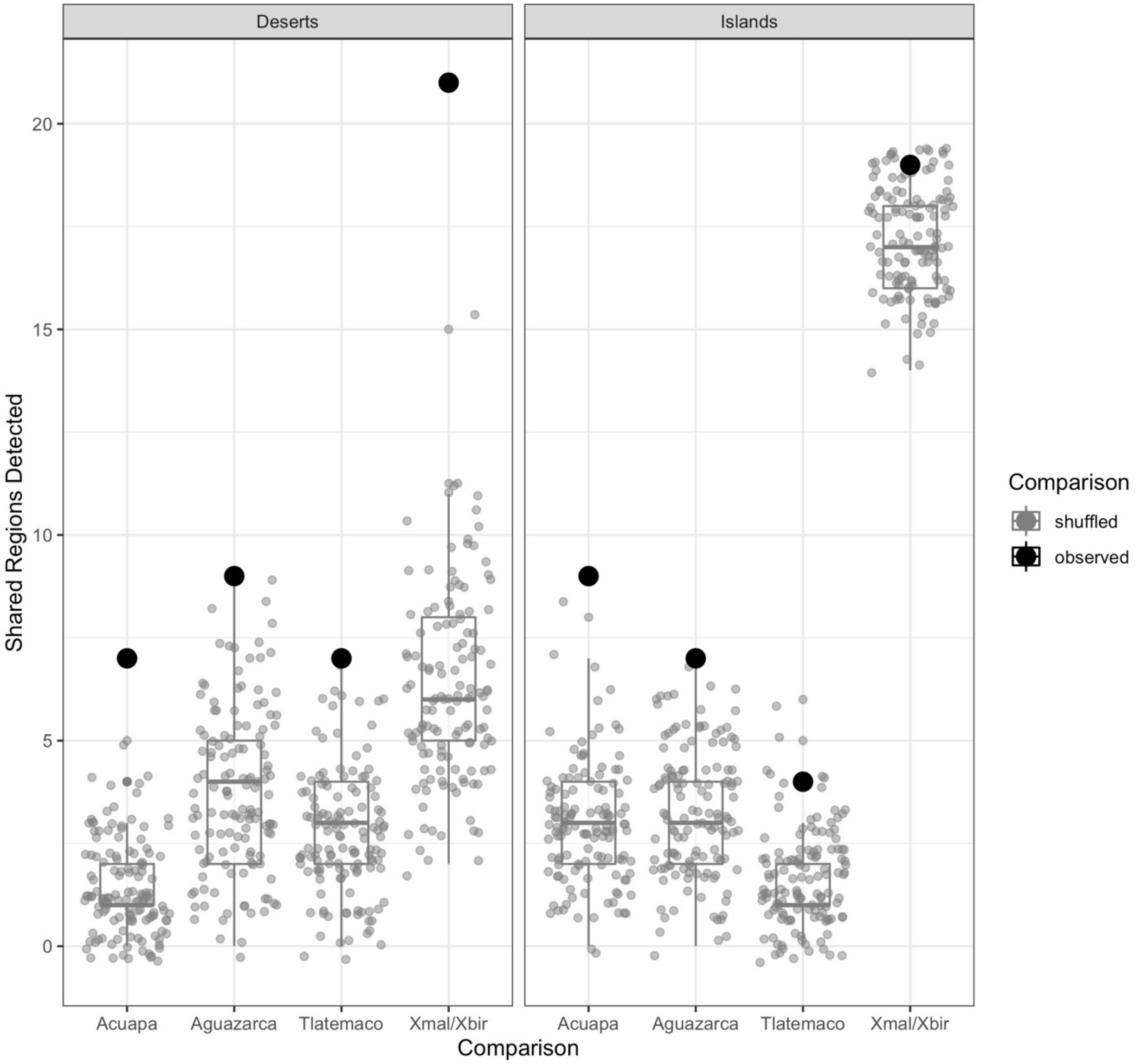


**Fig. S12.** Shared minor parent deserts between *X. birchmanni* × *X. cortezi* and *X. birchmanni* × *X. malinche* hybrid populations are enriched compared to expectations by chance when using a permutation approach to generate null datasets that preserves the structure of local ancestry correlation in the genome (see Supporting Information 8). By contrast, minor parent islands are less enriched compared to null datasets when using this approach. Results shown here indicate the number of shared minor parent deserts (or islands) between the Santa Cruz *X. birchmanni* × *X. cortezi* hybrid population and each *X. birchmanni* × *X. malinche* hybrid population (Acuapa, Aguazarca, and Tlatemaco). Large black circles show the observed number of shared minor parent deserts or islands. Gray points and boxplots show the expectations from 130 shuffled datasets tiling the genome (see Supporting Information 8). The column labeled Xmal/Xbir shows the number of shared deserts or islands between the Santa Cruz *X. birchmanni* × *X. cortezi* hybrid population and any single *X. birchmanni* × *X. malinche* population.

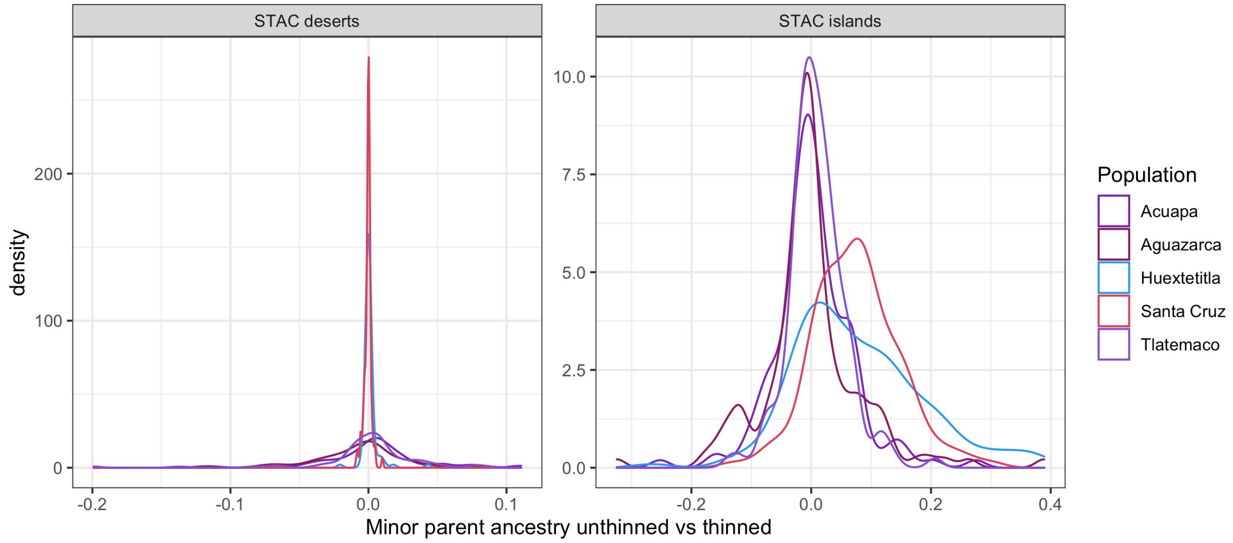


**Fig. S13**. We identified minor parent islands and deserts using genotypes from ancestry informative sites across the entire genome (referred to as the "unthinned" dataset in the main text). To ensure that minor parent islands and deserts were not generated as an artifact of variation in power to call ancestry along the genome, we re-calculated average ancestry in these regions using ancestry posterior probabilities generated from an input set of ancestry informative markers that were thinned to reduce power differences between different regions of the genome (see Methods; *Local ancestry inference in X. birchmanni* × *X. cortezi hybrids*). We found few differences in minor parent ancestry in deserts (**A**) based on this analysis. We identified more variation in ancestry in minor parent islands in the thinned data (**B**), and excluded a subset of these islands from further analysis (see Methods).

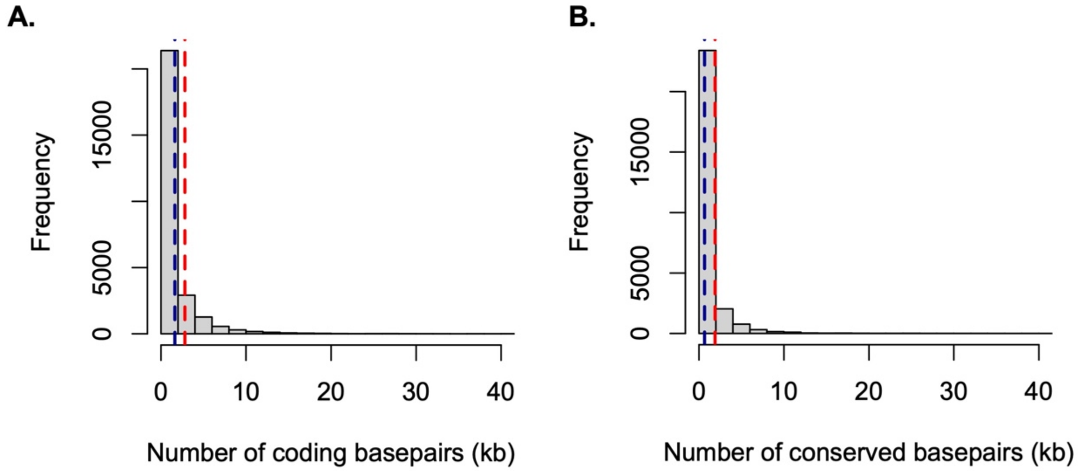


**Fig. S14.** Shared minor parent ancestry deserts and islands do not have an excess of coding (**A**) or conserved (**B**) basepairs compared to other regions of the genome that were not shared ancestry outliers. Gray distributions show number of coding and conserved basepairs in each 0.05 cM window across the genome. Red lines show the median number of coding or conserved basepairs in the 0.05 cM window that is the midpoint of the shared minor parent ancestry deserts. Blue lines show the median number of coding or conserved basepairs in the 0.05 cM window that is the midpoint of the shared minor parent ancestry islands.

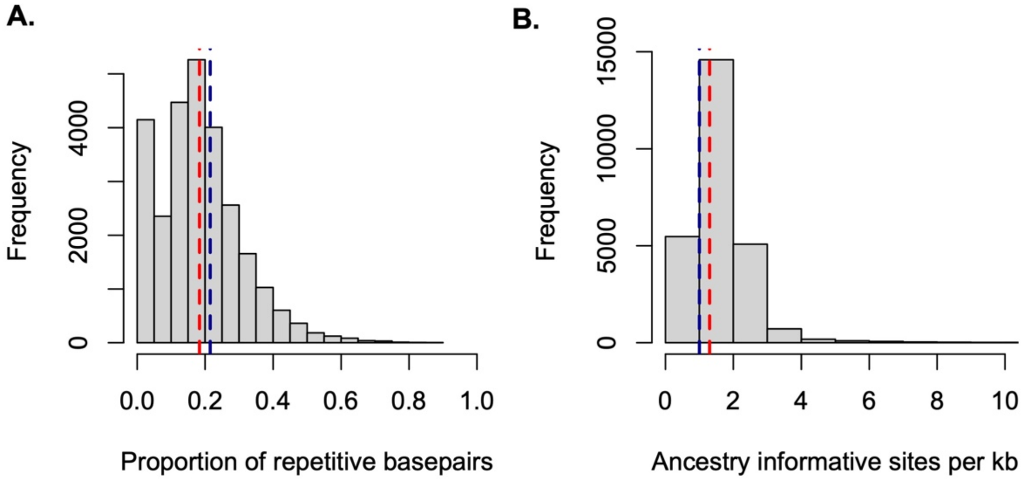


**Fig. S15.** Evaluation of density of repetitive elements (**A**) and ancestry informative sites (**B**) in minor parent ancestry deserts and islands relative to the genome-wide background. Distribution in gray shows 0.05 cM windows genome wide, red line shows the median value for the 0.05 cM window that is the midpoint of the shared minor parent ancestry desert, and blue line shows the median value for the 0.05 cM window that is the midpoint of the shared minor parent ancestry islands.

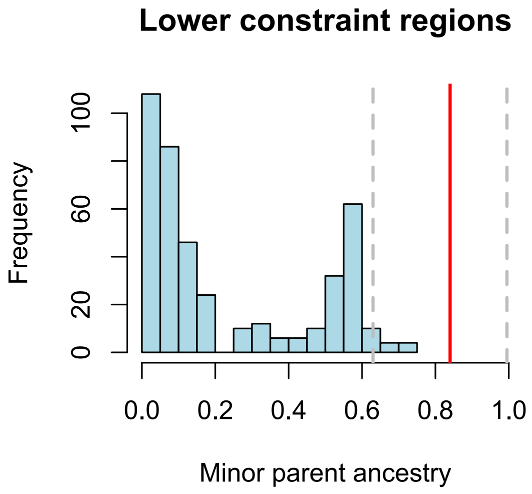


**Fig. S16.** Ancestry in minor parent islands compared to regions of low constraint. The blue distribution shows minor parent ancestry in the Santa Cruz population in 10 kb windows that are greater than 100 kb from the nearest coding basepair and with an inferred recombination rate in the upper 50% quantile of the genome-wide distribution. The red line shows the average minor parent ancestry in minor parent islands and the gray dashed lines shows the 95% confidence intervals. Thus, in addition to harboring a typical number of coding and conserved basepairs, minor parent islands are still ancestry outliers when compared to regions of the genome expected to have especially low constraint.

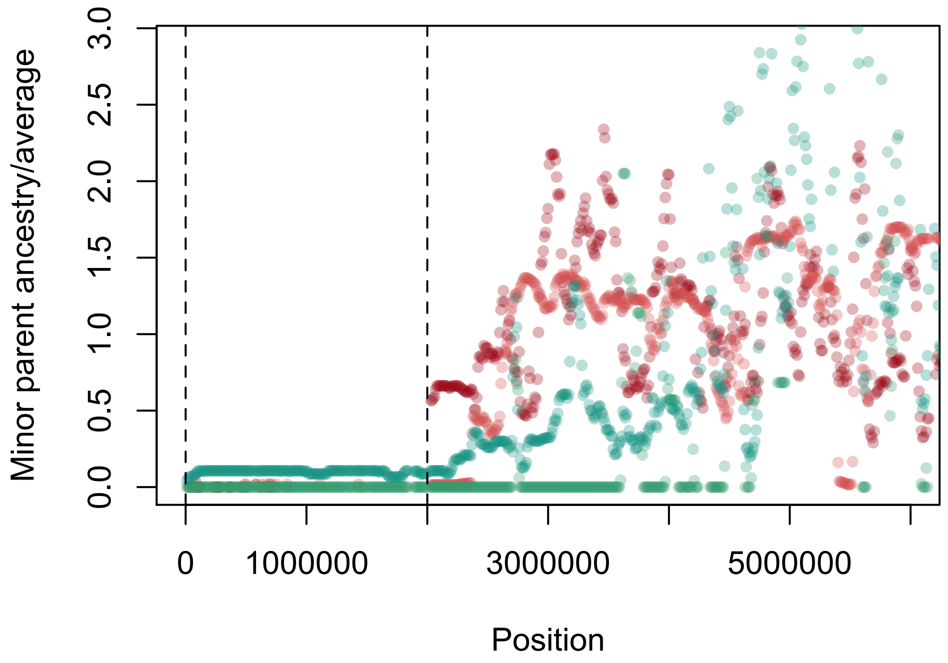


**Fig. S17.** Populations in which *X. birchmanni* is the minor parent have a large ~1 Mb desert of *X. birchmanni* ancestry on chromosome 21, the putative sex chromosome. Plotted here is minor parent ancestry in 10 kb windows relative to average minor parent ancestry genome-wide. Green indicates data from *X. birchmanni* × *X. cortezi* populations and red indicates data from *X. birchmanni* × *X. malinche* populations.

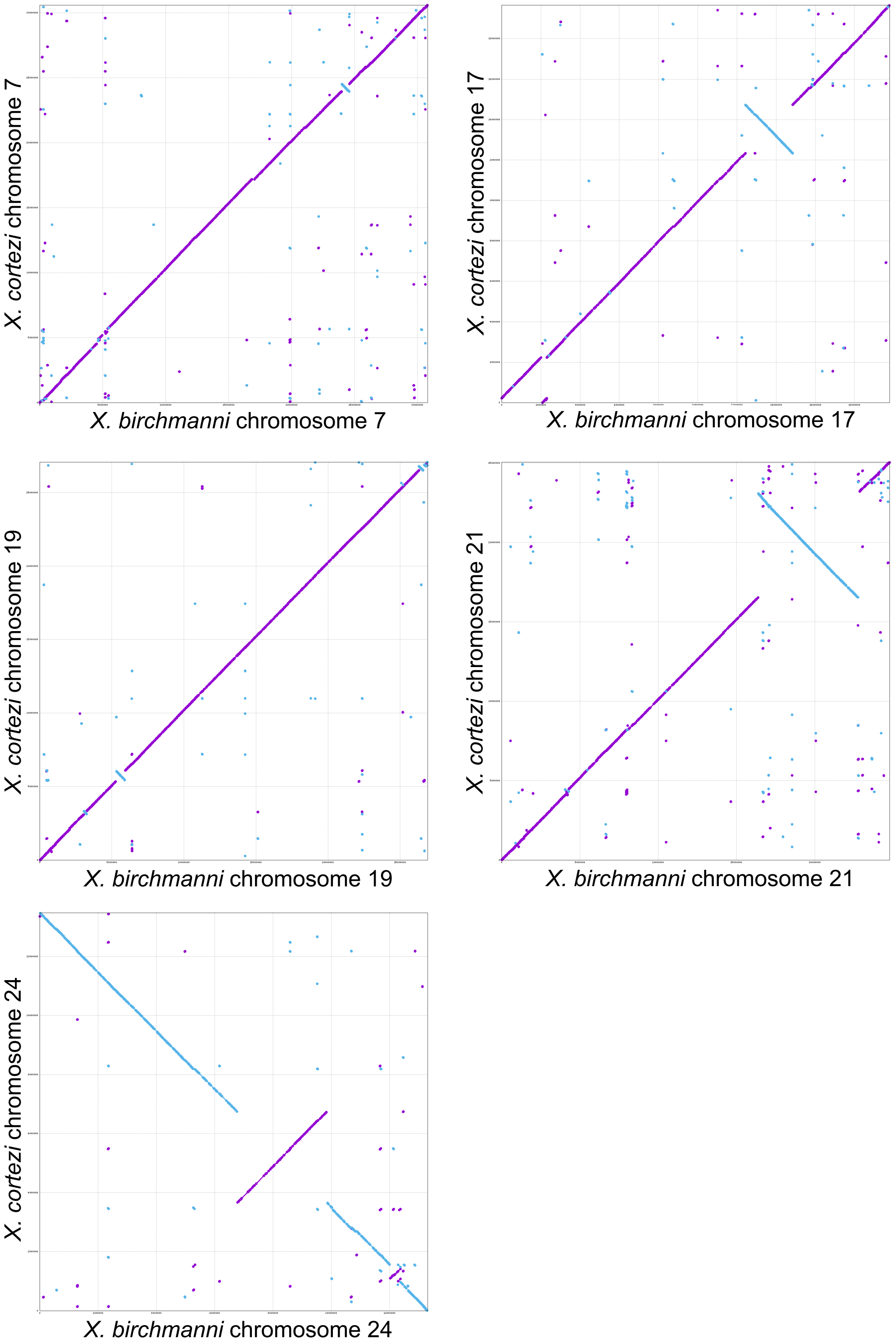


**Fig. S18.** MUMMER alignments indicate that the *X. birchmanni* and *X. cortezi* genomes are co-linear with the exception of several small inversions and the large inversions on chromosome 21 and chromosome 24 shown here. Alignments shown here also include chromosomes where a shared minor parent desert or island was found to overlap with an inversion.

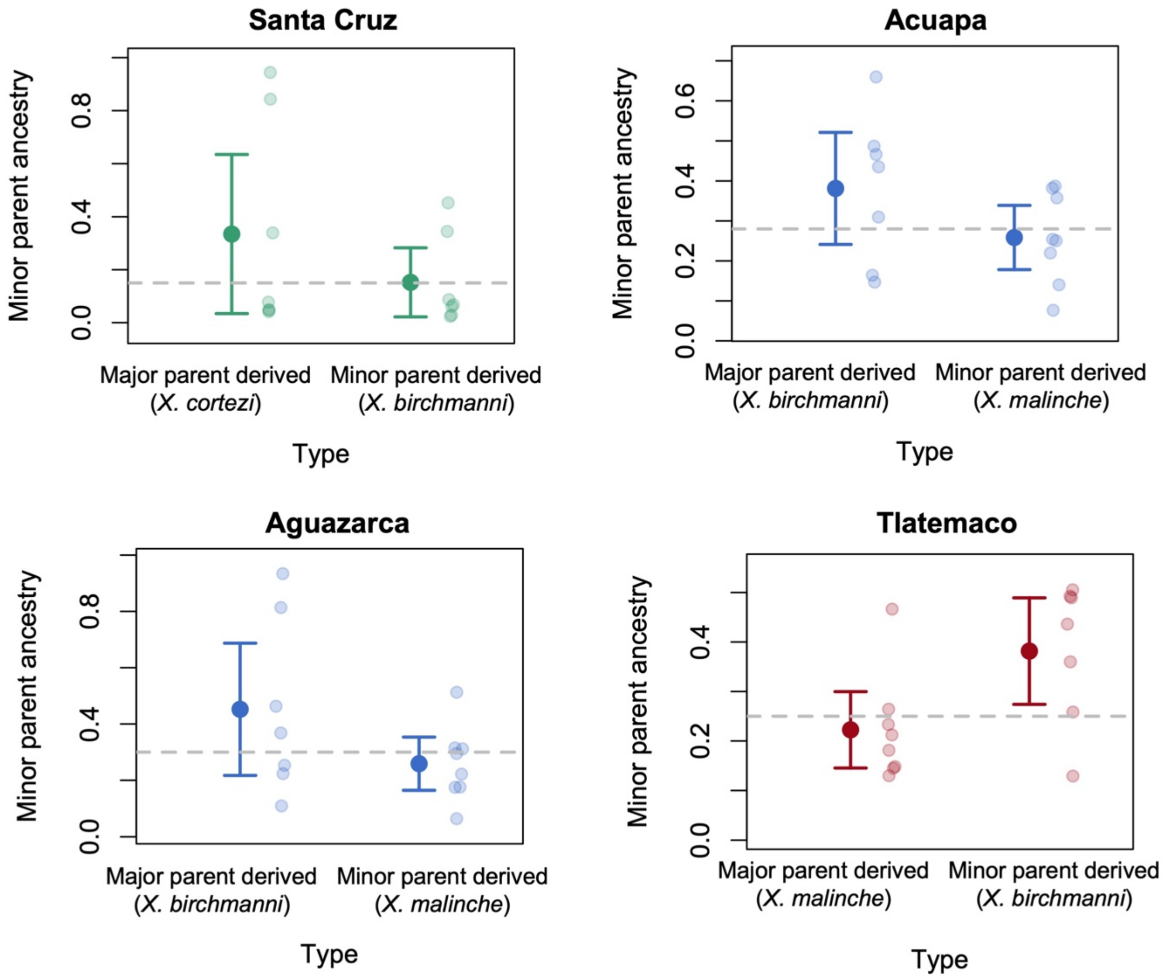


**Fig. S19.** Each parental species in our analysis differs from other species in a number of structural rearrangements and phylogenetic analysis allowed us to determine whether these rearrangements are likely derived in a particular species (see Methods). Here we plot minor parent ancestry at inversions that are derived in the major versus minor parent in each independent hybrid population. We find that inversions have unexpectedly high minor parent ancestry regardless of their origin in all hybrid populations, and in several populations inversions derived from the major parent are at unexpectedly low frequencies. Semi-transparent dots show ancestry at individual inversions, solid points and whiskers show the mean ancestry ± 2 standard errors of the mean. Gray line shows average minor parent ancestry in that population genome-wide.

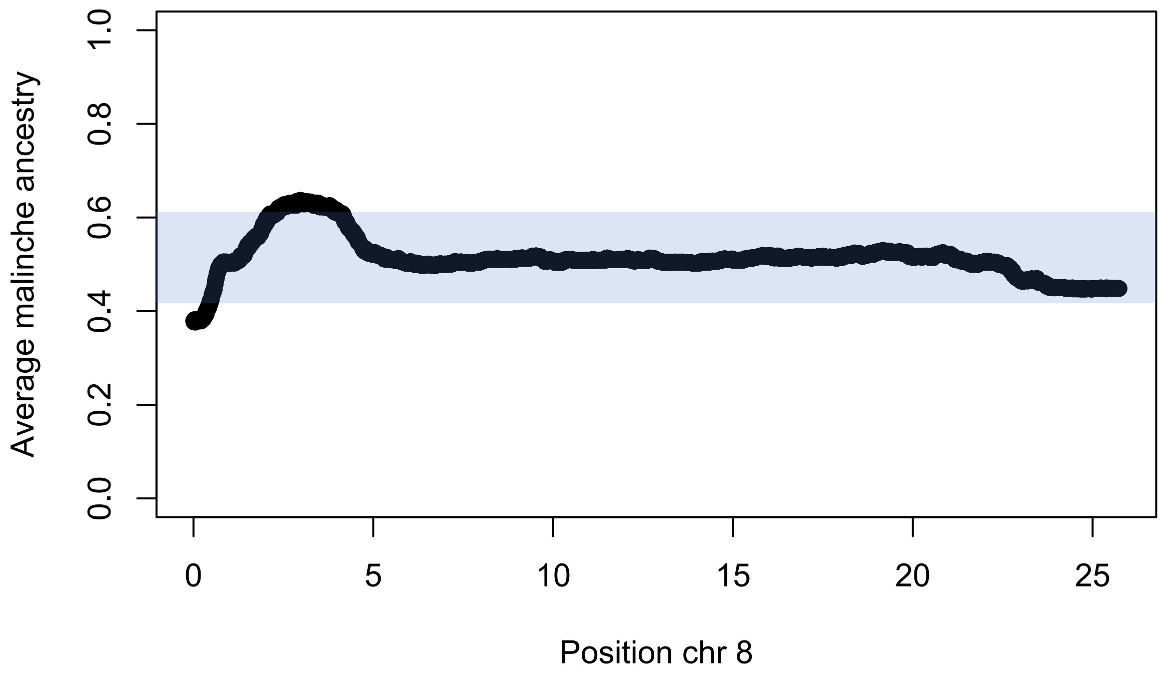


**Fig. S20.** Example of segregation distortion in $F_2$ hybrids between *X. birchmanni* and *X. malinche*. Given the cross design of an $F_1$ intercross we expect 50-50 segregation for parental ancestry types. Indeed, genome-wide average ancestry is 50.3% *X. malinche*. Plotted here is average ancestry by site along chromosome 8. Chromosome 8 has two regions that fall outside of the 99% confidence intervals for ancestry in the cross (shown by the blue shading).

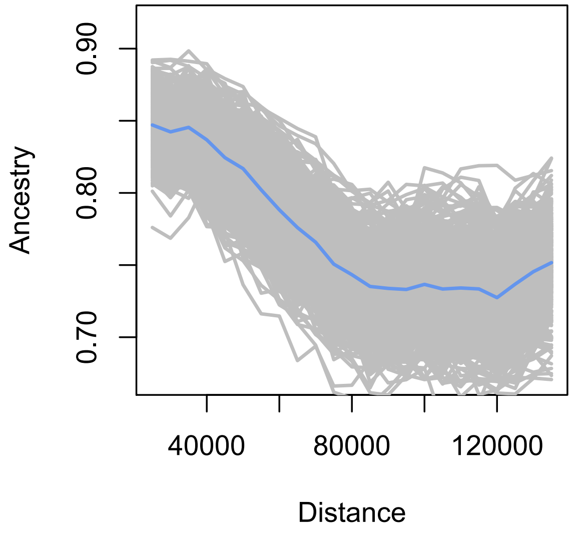


**Fig. S21.** *X. cortezi* ancestry in *X. birchmanni* × *X. cortezi* hybrid populations as a function of distance to segregation distorters identified in *X. birchmanni* × *X. malinche* early generation hybrids, excluding the segregation distorter on chromosome 6 that is also a shared ancestry desert with low minor parent ancestry across all hybrid populations.

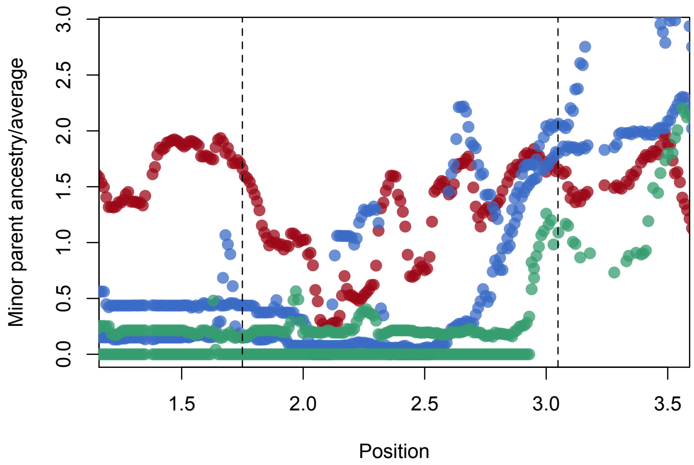


**Fig. S22.** Local ancestry summarized in 10 kb windows for all hybrid populations across the region on chromosome 13 associated with the shared ancestry desert on chromosome 6. Red – Tlatemaco (*malinche* × *birchmanni*), Blue – Acuapa and Aguazarca (*birchmanni* × *malinche*), Green –Santa Cruz (*birchmanni* × *cortezi*).

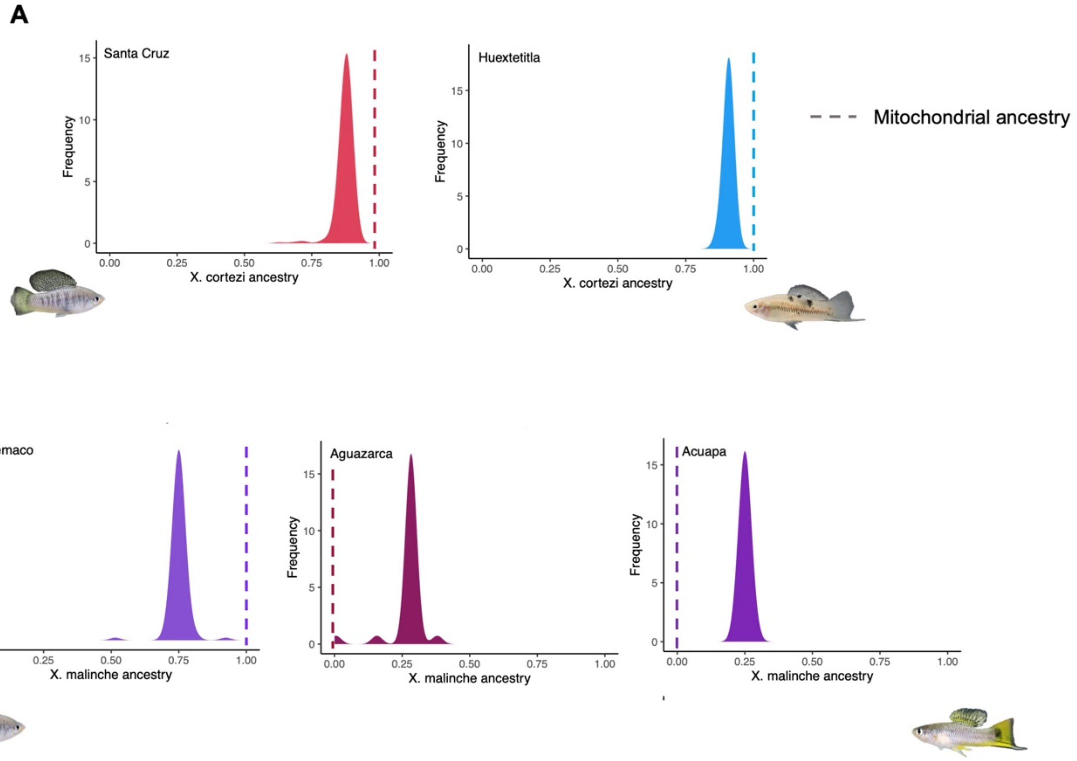


**Fig. S23**. All hybrid populations included in this study were fixed for the mitochondrial haplotype derived from the majority parental species. Shown here are the genome-wide ancestry distributions for individuals in *X. birchmanni* × *X. cortezi* hybrid populations (**A**) and *X. birchmanni* × *X. malinche* hybrid populations (**B**). Dotted lines show the ancestry of the mitochondrial haplotype for which the population is fixed.

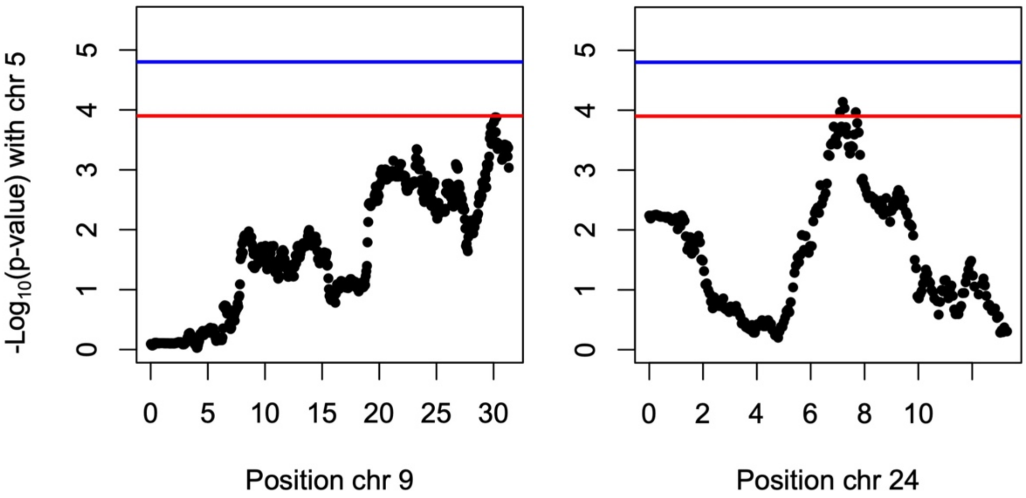


**Fig. S24.** Potential interactions between the chromosome 5 shared ancestry desert and two other regions of the genome (chromosome 9 and chromosome 24). Association between chromosome 5 desert and chromosome 9 and chromosome 24, detected at a FPR of 10% (red line). The FPR 5% threshold is also shown (blue line). Several known gene interactions exist between these three regions: chromosome 24 and chromosome 5 - *RBM43* and *trim25*, *rnd3b* and *rasal3*, chromosome 9 and chromosome 24 - *prrx1a* and *ccnt2b*.

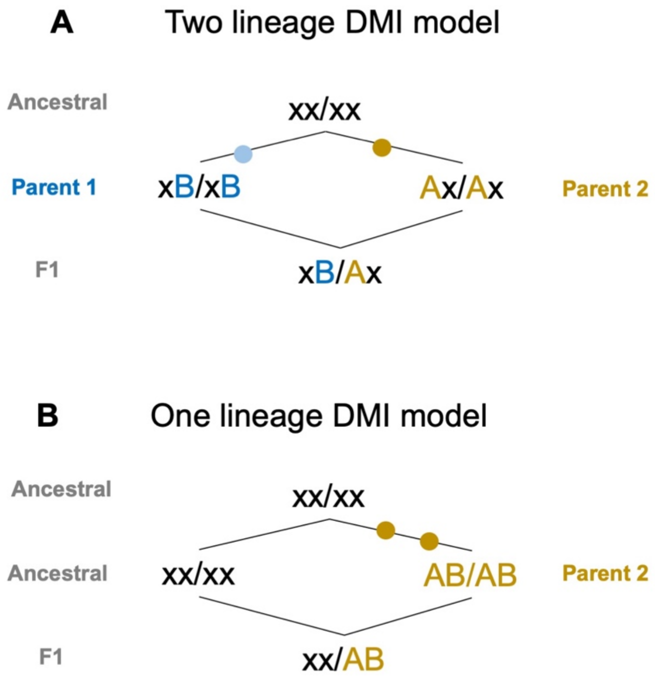


**Fig. S25.** Two possible routes through which Dobzhansky Muller Incompatibilities (DMIs) may arise between diverging lineages. **A**. As typically depicted, a derived mutation may arise in each lineage (A and B) which has the potential to negatively interact in hybrids. **B**. DMIs may also arise between the ancestral genotype (denoted as x alleles) and derived alleles that have accumulated on one lineage. This latter scenario may be a possible route through which shared hybrid incompatibilities accumulate between related species, if one lineage has fixed several substitutions and others retain the ancestral genotype.

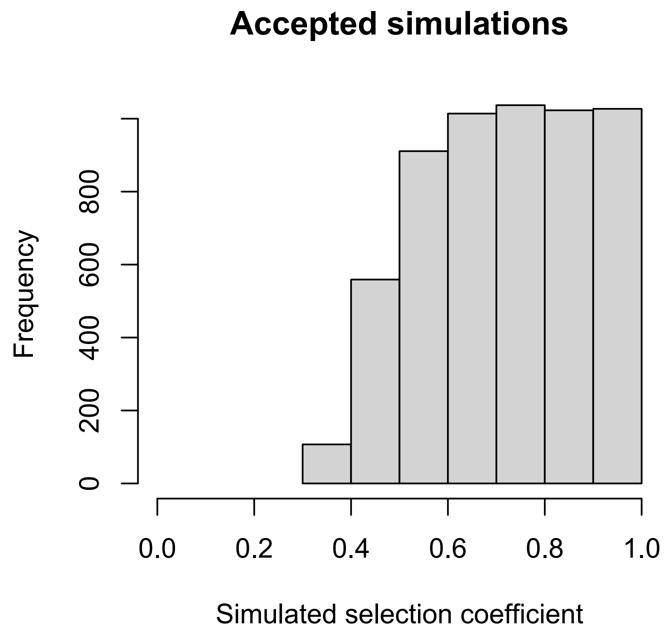


**Fig. S26.** In the main text, we identify segregation distorters based on local ancestry data from $F_2$ hybrids generated between *X. birchmanni* × *X. malinche* that deviate from the expected 50-50 ancestry frequency at a given locus. We performed simulations to ask what selection coefficients are consistent with the deviations from expected admixture proportions that we observe at segregation distortion loci. Shown here is the distribution of accepted selection coefficients from simulations (prior *s* 0-1); see Methods for simulation descriptions.

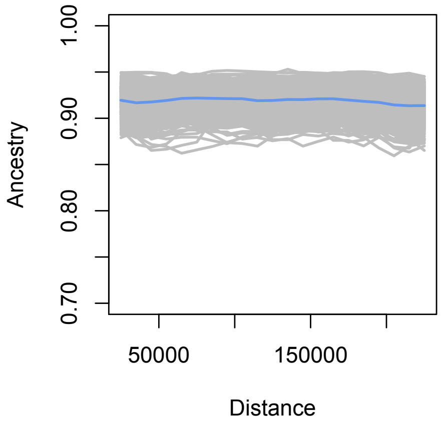


**Fig. S27.** Although we see high *X. cortezi* ancestry in Huextetitla and Santa Cruz near sites that are strongly selected in *X. birchmanni* × *X. malinche* hybrids (Fig. 3), we do not see a signature of higher *X. cortezi* ancestry near putative segregating DMIs identified in natural *X. birchmanni* × *X. malinche* hybrid populations. Shown here are the results for the Santa Cruz population as a function of distance to these sites. Gray lines show results of 500 replicates bootstrap resampling the data, blue shows the average across simulations.

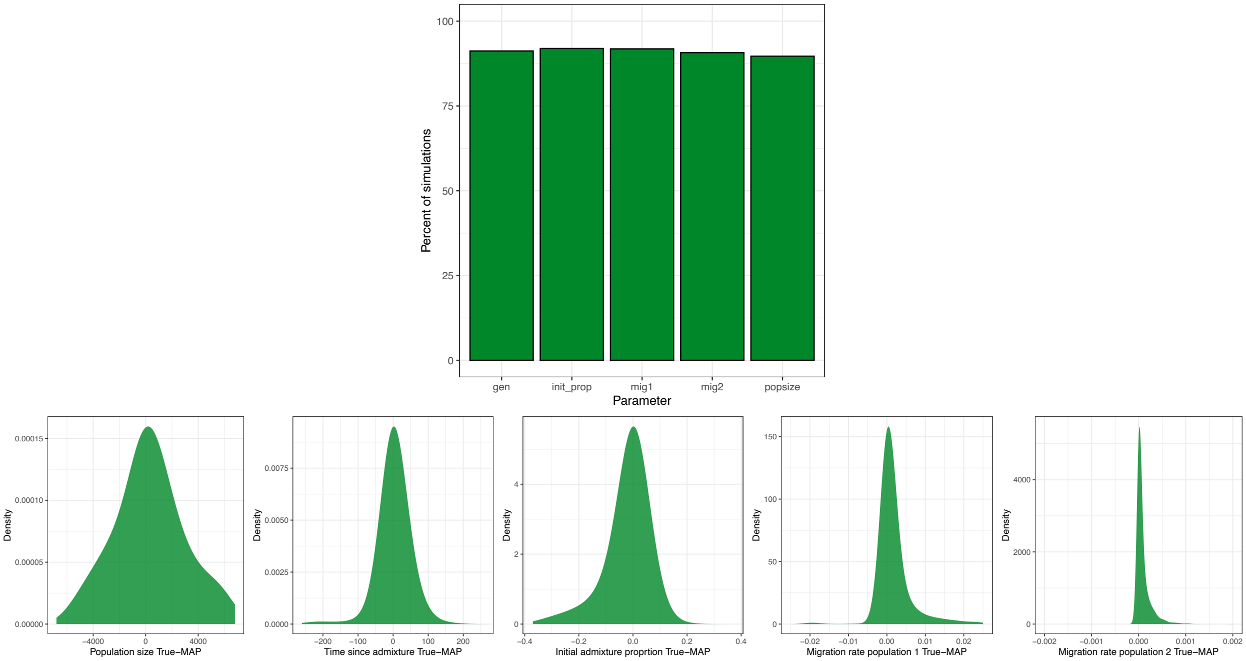


**Fig. S28.** Proof of principle simulations for Approximate Bayesian Computation (ABC) approach. When we perform inference on known simulations, we recover the true input parameter within the 95% inner quantile of the posterior distribution for >90% of the (5,000) simulations we tested (top). The difference between the true input value and the mode of the posterior for all parameters centers on zero (bottom). We accepted simulations based on three criteria: median length of minor parent ancestry tracts, mean hybrid index, and coefficient of variation in ancestry chromosome-wide across individuals. For the purpose of this proof of principle test, we required that randomly sampled parameter sets and corresponding summary statistics resulted in >500 accepted simulations and randomly sampled down to 500 accepted simulations. See Supporting Information 3 for a full description of these simulations.

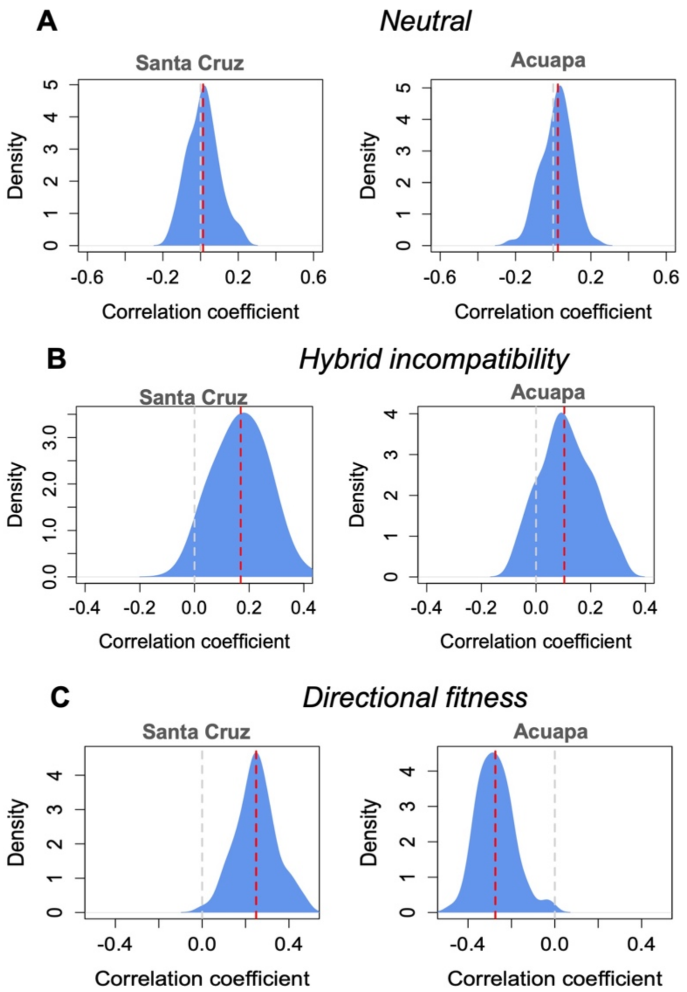


**Fig. S29.** Summary of within population correlations between minor parent ancestry and recombination rate across 100 replicate simulations of different scenarios of selection on hybrids. **A**. In neutral simulations matching the inferred demographic history of Santa Cruz and Acuapa, average correlation coefficients between minor parent ancestry and recombination rate (red line) fall close to the expected value of zero (gray line). **B**. In models implementing incompatibility selection against hybrids, average correlation coefficients between minor parent ancestry and recombination rate (red line) are positive. **C**. In contrast, in models where there is directional selection in favor of ancestry from one parent species (in these simulations, *X. cortezi* ancestry), simulations predict that the direction of the correlations between minor parent ancestry and recombination rate will vary depending on the population's admixture proportion.

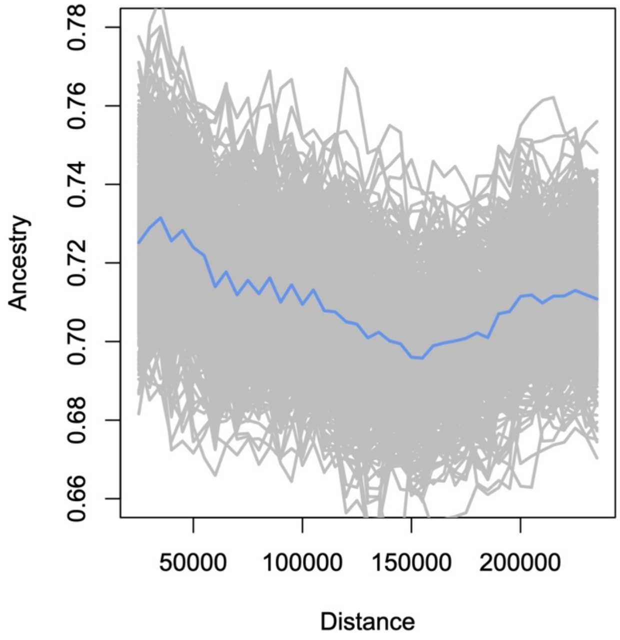


**Fig. S30.** Average major parent ancestry in Tlatemaco, a natural *X. birchmanni* x *X. malinche* hybrid population, as a function of the distance to the closest segregation distorter. Segregation distorters were identified in $F_2$ crosses between *X. birchmanni* and *X. malinche*. These results suggest that minor parent ancestry tends to be higher at segregation distorters in the Tlatemaco population, despite weak correlations in local ancestry genome-wide between Tlatemaco and other hybrid populations. The blue line shows average major parent ancestry, gray lines show the results from 500 replicates of bootstrap resampling the data.

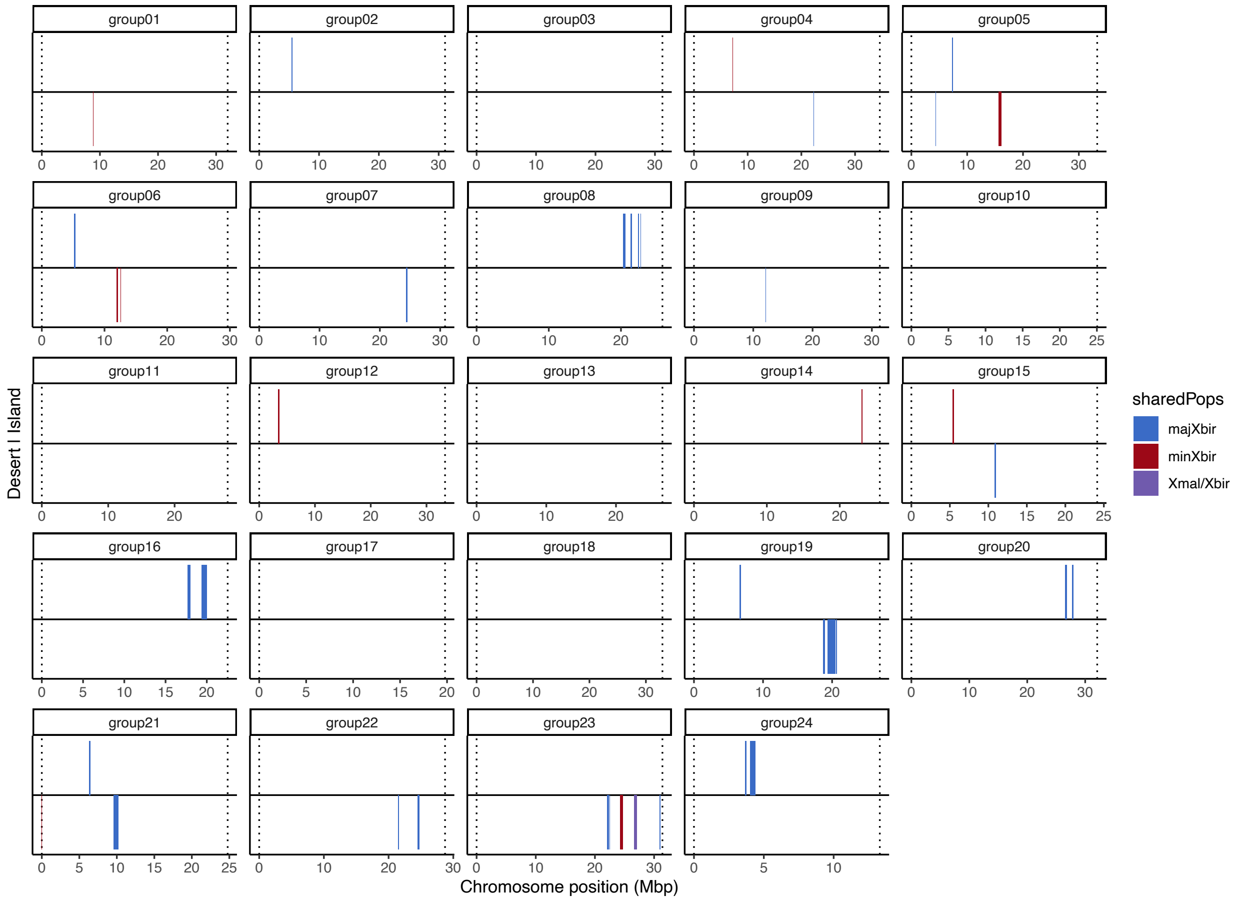


**Fig. S31.** Locations of minor parent deserts and islands that were detected in the Santa Cruz population and are shared in at least one *X. birchmanni* × *X. malinche* replicate population. Locations are shown based on their position across the *X. birchmanni* reference genome. In each plot any colored bar above the horizontal line is an island and any below is a desert.

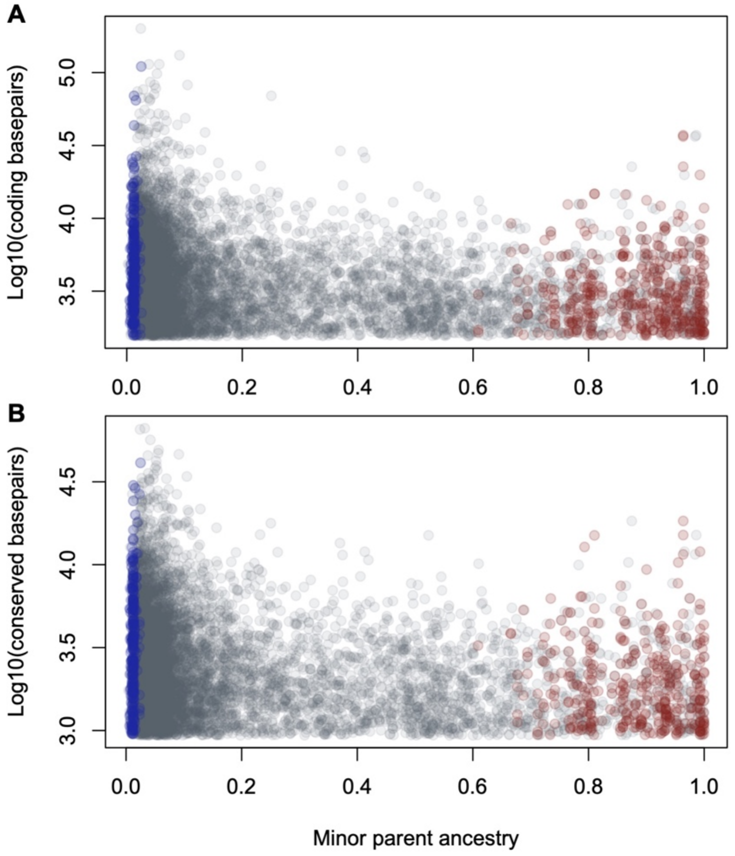


**Fig. S32.** Minor parent deserts (blue) and islands (red) are still ancestry outliers when compared only to 0.05 cM windows (gray) with a similar density of coding (**A**) or conserved basepairs (**B**). Shown here are minor parent deserts and islands that fall in the top 25% quantile in terms of the number of linked coding or conserved basepairs, versus all 0.05 cM windows genome-wide that fall into this same coding or conserved quantile (gray).

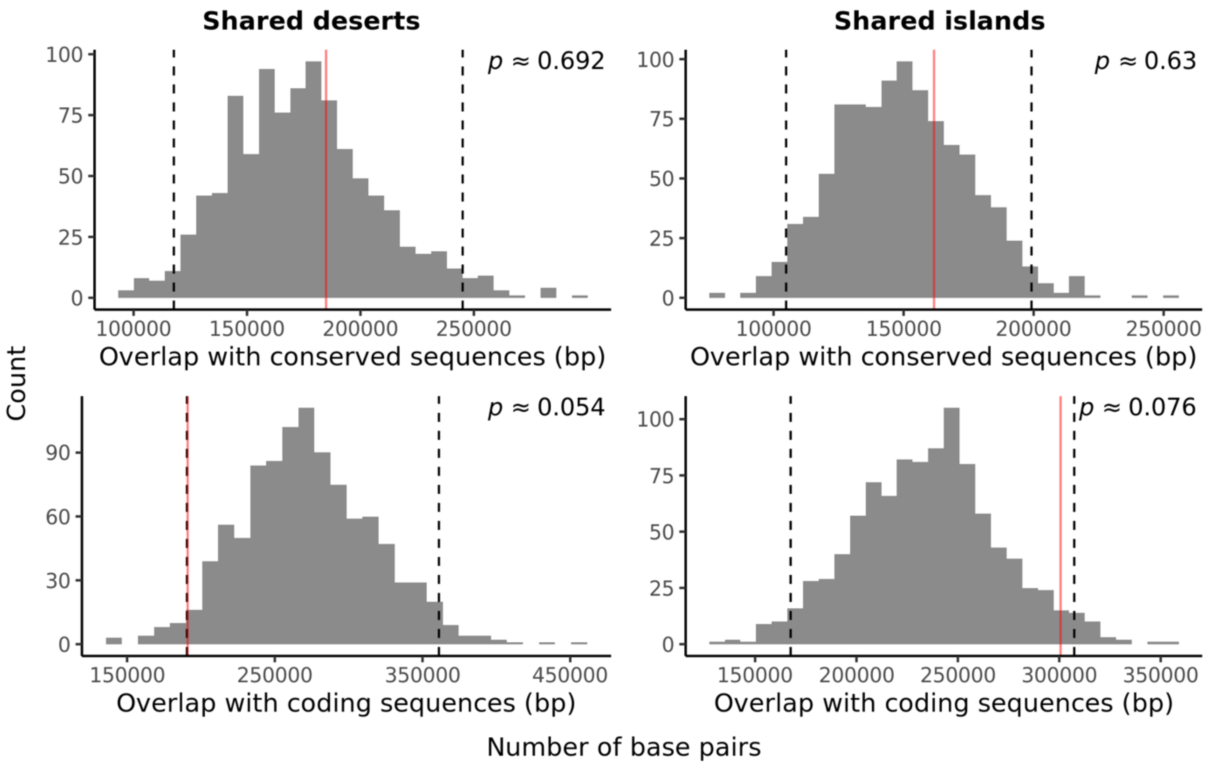


**Fig. S33.** Overlap of conserved (top row) and coding (bottom row) base pairs in shared deserts and islands versus matched null permutations. Red lines represent the observed values in shared deserts and islands. Dashed lines represent 2.5% and 97.5% quantiles; *p*-values were estimated empirically based on comparisons of the focal and the permutated data.

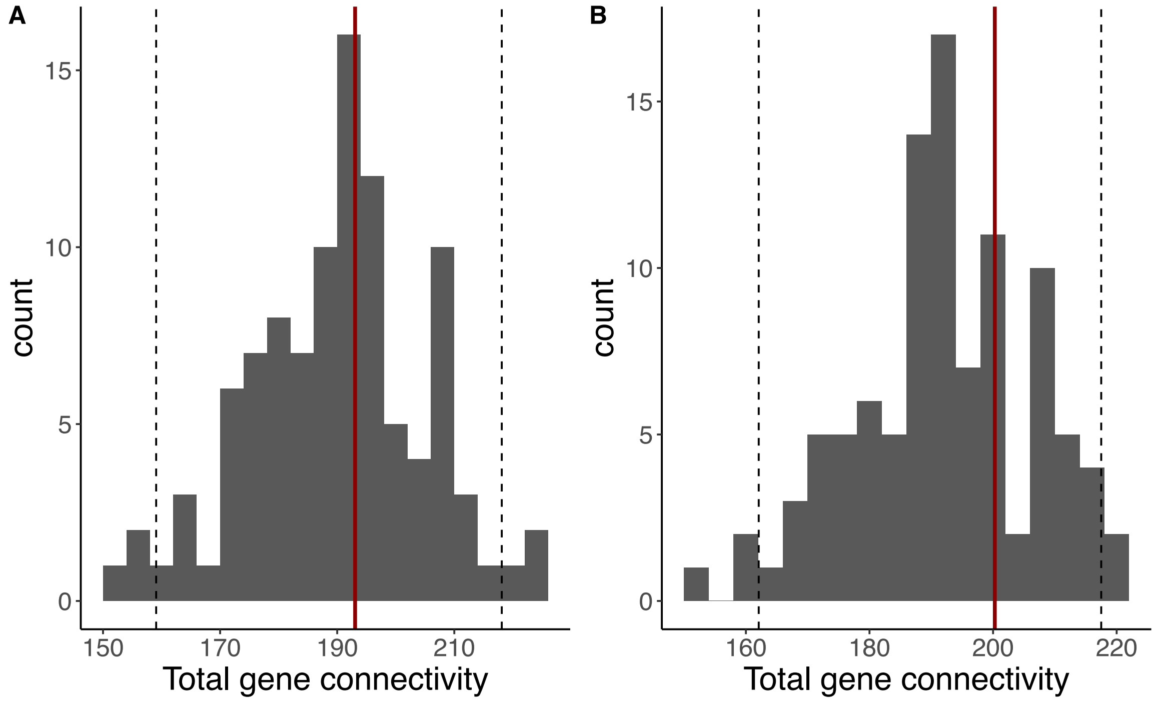


**Fig. S34.** Mean total connectivity based on WGCNA analysis of genes in shared minor parent (**A**) deserts and (**B**) islands, marked by the red line, compared to the distribution of mean total connectivity for matched null gene sets, shown by the grey distribution. The black dashed lines show the 95% confidence intervals for each null distribution.

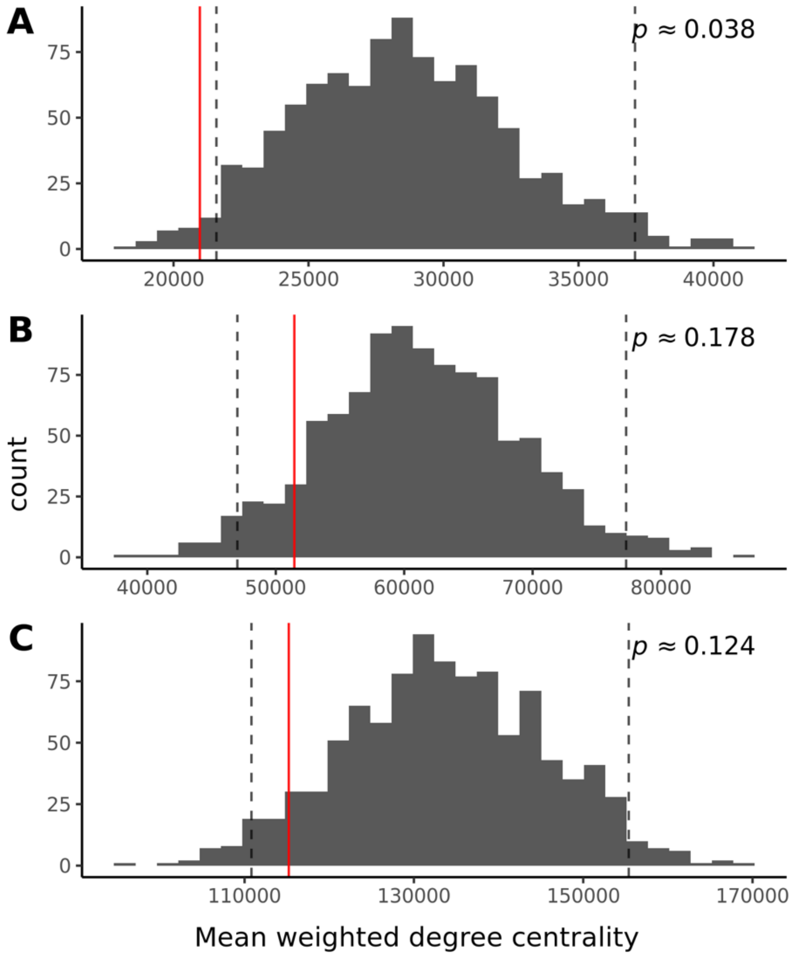

**Fig. S35.** Average STRING-db connectivity of genes in shared minor parent deserts (red line) compared to genes in 1,000 matched null permutations based on (**A**) co-expression, (**B**) experiments/biochemistry, and (**C**) combined interaction score categories. Dashed lines represent 2.5% and 97.5% quantiles; *p*-values were estimated empirically based on comparisons of the focal and the permutated data.

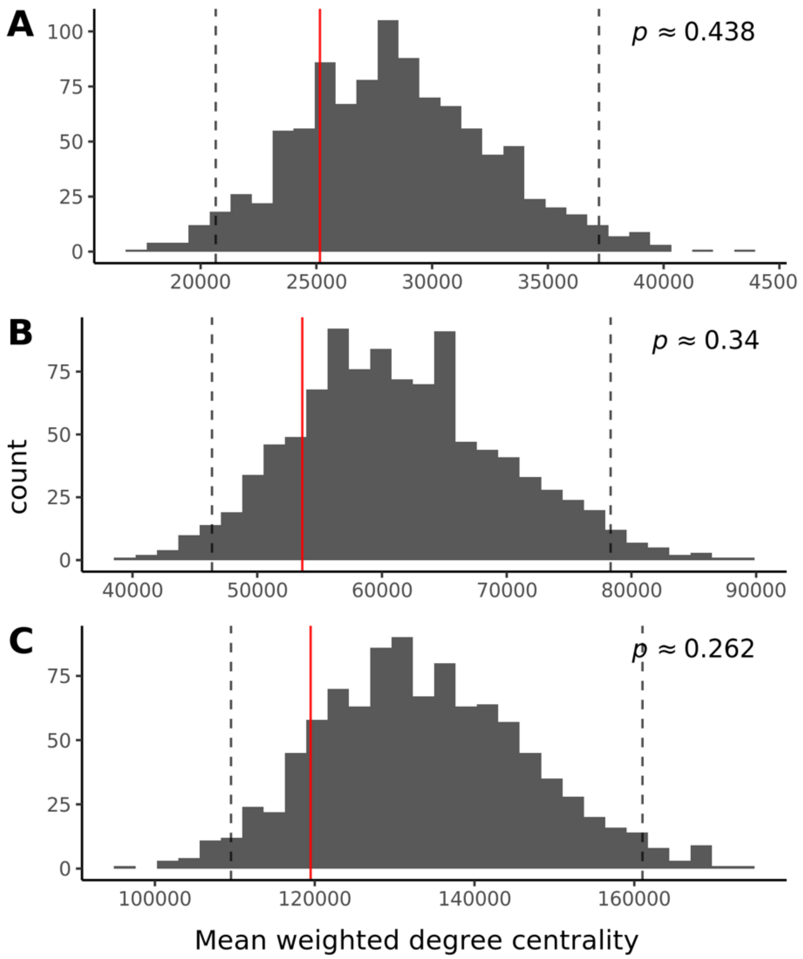


**Fig. S36.** Average STRING-db connectivity of genes in shared minor parent islands (red line) compared to genes in 1,000 matched null permutations based on (**A**) co-expression, (**B**) experiments/biochemistry, and (**C**) combined interaction score categories. Dashed lines represent 2.5% and 97.5% quantiles; *p*-values were estimated empirically based on comparisons of the focal and the permutated data.

### Supporting Information References


1. Culumber ZW, Fisher HS, Tobler M, Mateos M, Barber PH, Sorenson MD, et al. Replicated hybrid zones of Xiphophorus swordtails along an elevational gradient. Molecular Ecology. 2011;20: 342–356. doi:10.1111/j.1365-294X.2010.04949.x

2. Culumber ZW, Shepard DB, Coleman SW, Rosenthal GG, Tobler M. Physiological adaptation along environmental gradients and replicated hybrid zone structure in swordtails (Teleostei: Xiphophorus). J Evol Biol. 2012;25: 1800–1814. doi:10.1111/j.1420-9101.2012.02562.x

3. Schumer M, Powell DL, Delclós PJ, Squire M, Cui R, Andolfatto P, et al. Assortative mating and persistent reproductive isolation in hybrids. Proc Natl Acad Sci USA. 2017;114: 10936. doi:10.1073/pnas.1711238114

4. Culumber ZW, Tobler M. Ecological divergence and conservatism: spatiotemporal patterns of niche evolution in a genus of livebearing fishes (Poeciliidae: Xiphophorus). BMC Evol Biol. 2016;16: 44. doi:10.1186/s12862-016-0593-4

5. Powell DL, Moran B, Kim B, Banerjee SM, Aguillon SM, Fascinetto-Zago P, et al. Two new hybrid zones expand the swordtail hybridization model system. bioRxiv. 2020; 2020.11.18.389205. doi:10.1101/2020.11.18.389205

6. Li H, Handsaker B, Wysoker A, Fennell T, Ruan J, Homer N, et al. The Sequence Alignment/Map format and SAMtools. Bioinformatics. 2009;25: 2078–2079. doi:10.1093/bioinformatics/btp352

7. Li H. A statistical framework for SNP calling, mutation discovery, association mapping and population genetical parameter estimation from sequencing data. Bioinformatics. 2011;27: 2987–2993. doi:10.1093/bioinformatics/btr509

8. Purcell S, Neale B, Todd-Brown K, Thomas L, Ferreira MAR, Bender D, et al. PLINK: A Tool Set for Whole-Genome Association and Population-Based Linkage Analyses. The American Journal of Human Genetics. 2007;81: 559–575. doi:10.1086/519795

9. Cui R, Schumer M, Rosenthal GG. Admix'em: a flexible framework for forward-time simulations of hybrid populations with selection and mate choice. Bioinformatics. 2016;32: 1103–1105. doi:10.1093/bioinformatics/btv700

10. Schumer M, Cui R, Powell DL, Dresner R, Rosenthal GG, Andolfatto P. High-resolution mapping reveals hundreds of genetic incompatibilities in hybridizing fish species. McVean G, editor. eLife. 2014;3: e02535. doi:10.7554/eLife.02535

11. Powell DL, García-Olazábal M, Keegan M, Reilly P, Du K, Díaz-Loyo AP, et al. Natural hybridization reveals incompatible alleles that cause melanoma in swordtail fish. Science. 2020;368: 731–736. doi:10.1126/science.aba5216

12. Lu Y, Sandoval A, Voss S, Lai Z, Kneitz S, Boswell W, et al. Oncogenic allelic interaction in Xiphophorus highlights hybrid incompatibility. PNAS. 2020;117: 29786–29794. doi:10.1073/pnas.2010133117

13. Haller BC, Messer PW. SLiM 3: Forward Genetic Simulations Beyond the Wright–Fisher Model. Hernandez R, editor. Molecular Biology and Evolution. 2019;36: 632–637. doi:10.1093/molbev/msy228

14. Haller BC, Galloway J, Kelleher J, Messer PW, Ralph PL. Tree-sequence recording in SLiM opens new horizons for forward-time simulation of whole genomes. bioRxiv. 2018; 407783. doi:10.1101/407783

15. Schumer M, Xu C, Powell DL, Durvasula A, Skov L, Holland C, et al. Natural selection interacts with recombination to shape the evolution of hybrid genomes. Science. 2018;360: 656. doi:10.1126/science.aar3684

16. Baker Z, Schumer M, Haba Y, Bashkirova L, Holland C, Rosenthal GG, et al. Repeated losses of PRDM9-directed recombination despite the conservation of PRDM9 across vertebrates. In: eLife [Internet]. 6 Jun 2017 [cited 23 Jul 2019]. doi:10.7554/eLife.24133

17. Singhal S, Leffler EM, Sannareddy K, Turner I, Venn O, Hooper DM, et al. Stable recombination hotspots in birds. Science. 2015;350: 928–932. doi:10.1126/science.aad0843

18. Lam I, Keeney S. Nonparadoxical evolutionary stability of the recombination initiation landscape in yeast. Science. 2015;350: 932–937. doi:10.1126/science.aad0814

19. Siepel A, Bejerano G, Pedersen JS, Hinrichs AS, Hou M, Rosenbloom K, et al. Evolutionarily conserved elements in vertebrate, insect, worm, and yeast genomes. Genome Res. 2005;15: 1034–1050. doi:10.1101/gr.3715005

20. Telis N, Aguilar R, Harris K. Selection against archaic hominin genetic variation in regulatory regions. Nature Ecology & Evolution. 2020; 1–9. doi:10.1038/s41559-020-01284-0

21. Moran BM, Payne C, Langdon Q, Powell DL, Brandvain Y, Schumer M. The genetic consequences of hybridization. arXiv:201204077 [q-bio]. 2020 [cited 27 Feb 2021]. Available: http://arxiv.org/abs/2012.04077

22. Juric I, Aeschbacher S, Coop G. The Strength of Selection against Neanderthal Introgression. PLOS Genetics. 2016;12: e1006340. doi:10.1371/journal.pgen.1006340

23. Hahn MW. Accurate Inference and Estimation in Population Genomics. Molecular Biology and Evolution. 2006;23: 911–918. doi:10.1093/molbev/msj094

24. Quinlan AR, Hall IM. BEDTools: a flexible suite of utilities for comparing genomic features. Bioinformatics. 2010;26: 841–842. doi:10.1093/bioinformatics/btq033

25. Powell DL, Payne C, Banerjee SM, Keegan M, Bashkirova E, Cui R, et al. The Genetic Architecture of Variation in the Sexually Selected Sword Ornament and Its Evolution in Hybrid Populations. Current Biology. 2021 [cited 28 Jan 2021]. doi:10.1016/j.cub.2020.12.049

26. Coyne JA, Orr HA. Speciation. Sunderland, MA: Sinaeur Associates; 2004. Available: <Go to ISI>://ZOOREC:ZOOR14109056022

27. Schumer M, Cui R, Rosenthal GG, Andolfatto P. Reproductive Isolation of Hybrid Populations Driven by Genetic Incompatibilities. PLOS Genetics. 2015;11: e1005041. doi:10.1371/journal.pgen.1005041

28. Armstrong J, Hickey G, Diekhans M, Deran A, Fang Q, Xie D, et al. Progressive alignment with Cactus: a multiple-genome aligner for the thousand-genome era. bioRxiv. 2019; 730531. doi:10.1101/730531

29. Mack KL, Nachman MW. Gene Regulation and Speciation. Trends in Genetics. 2017;33: 68–80. doi:10.1016/j.tig.2016.11.003

30. Porter AH, Johnson NA. Speciation despite gene flow when developmental pathways evolve. Evolution. 2002;56: 2103–2111. doi:10.1111/j.0014-3820.2002.tb00136.x

31. Langfelder P, Horvath S. WGCNA: an R package for weighted correlation network analysis. BMC Bioinformatics. 2008;9: 559. doi:10.1186/1471-2105-9-559

32. Love MI, Huber W, Anders S. Moderated estimation of fold change and dispersion for RNA-seq data with DESeq2. Genome Biology. 2014;15: 550. doi:10.1186/s13059-014-0550-8

33. Szklarczyk D, Morris JH, Cook H, Kuhn M, Wyder S, Simonovic M, et al. The STRING database in 2017: quality-controlled protein–protein association networks, made broadly accessible. Nucleic Acids Res. 2017;45: D362–D368. doi:10.1093/nar/gkw937

34. Proceedings of the Python in Science Conference (SciPy): Exploring Network Structure, Dynamics, and Function using NetworkX. [cited 13 May 2021]. Available: http://conference.scipy.org/proceedings/SciPy2008/paper_2/

35. Gravel S. Population Genetics Models of Local Ancestry. Genetics. 2012;191: 607. doi:10.1534/genetics.112.139808
